# The Cardiovascular Benefit of Menopausal Hormone Treatment is Age-Dependent in *Ldlr^−/−^* Mice

**DOI:** 10.64898/2026.06.29.734522

**Authors:** Cassandra D. Lalisan, Connor Gavin, Bridget Litts, Min Zhong, Jeffrey A. Rein, Julia An, Rajguru Boobalan, Yu Wang, Amelia Van Aelst, Sivaprakasam Chinnarasu, Yaomin Xu, MacRae F. Linton, John M. Stafford, Lin Zhu

**Author notes:** Address for Correspondence: Lin Zhu, MD, PhD, The Ohio State University Wexner Medical Center Pelotonia Research Center, 2255 Kenny Road, Rm 5115, Columbus, OH 43210. These authors equally contribute to this manuscript (Cassandra D. Lalisan, Connor Gavin).

## Abstract

**Background:** The risk of atherosclerotic cardiovascular disease (ASCVD) increases in women after menopause, coinciding with declining estrogen levels. However, menopausal hormone therapy (MHT) has not consistently reduced ASCVD events in post-menopausal women, despite improvements in lipid profiles. The mechanisms underlying the diminished cardiovascular protective effects of estrogens during MHT remain poorly understood.

**Methods:** Atherosclerosis was established in 11-month-old female *Ldlr^−/−^* mice through Western diet feeding. To investigate the effects of menopause and 17β-estradiol (E_2_) treatment on atherosclerotic burden during lipid normalization, mice were switched to a chow diet and subjected to ovariectomy (OVX) with and without E_2_ treatment. To determine the contribution of inflammation to ASCVD risk during MHT, a cohort of mice received *Ifnγ^−/−^* bone marrow transplantation (*Ifnγ^−/−^*^BM^) before hormone treatment. To assess the influence of age on ASCVD risk during MHT, parallel studies were conducted in young mice, beginning at 12 weeks of age. Outcomes included lipid profiles, HDL function, circulating cytokines, atherosclerotic burden, liver lipid metabolic pathways and transcriptomic profiles, and hepatic estrogen receptor signaling.

**Results:** OVX increased atherosclerotic burden compared with sham-operation. E_2_ treatment following OVX failed to reduce atherosclerosis in older mice but reduced atherosclerosis in young mice during chow diet-induced lipid normalization. In older mice, E_2_ treatment elevated circulating IFNγ, impaired HDL antioxidant and cholesterol efflux functions, and increased plaque inflammation and features of plaque vulnerability. Suppression of inflammation through *Ifnγ^−/−^*^BM^ transplantation restored HDL function and the ability of E_2_ treatment to reduce atherosclerosis in older mice. In contrast, E_2_ treatment following OVX in young mice limited inflammation and improved HDL function. E_2_ treatment increased liver fatty acid metabolic signaling in both older and young mice but increased liver 4-hydroxynonenal (4-HNE) levels only in older mice. The age-dependent response was associated with remodeling of hepatic estrogen receptor (ER) expression, characterized by reduced ERα and increased G protein-coupled ER expression with aging. Liver transcriptomic analyses identified inflammatory and interferon-associated signatures following OVX+E_2_ treatment in older mice. qPCR analyses across older and young mice suggested age-associated differences in inflammatory transcripts in response to E_2_.

**Conclusions:** These findings suggest that aging alters hepatic responses to E_2_ treatment following ovariectomy, promoting oxidative stress and inflammatory signaling that contribute to persistent systemic inflammation and compromise the ability of E_2_ to reduce atherosclerosis.

## Introduction

Atherosclerotic cardiovascular disease (ASCVD) is a leading cause of mortality for women worldwide. Premenopausal women have substantially lower risk for metabolic disorders and typically experience ASCVD onset nearly a decade later than age-matched men^1,2^. This woman-specific cardiometabolic protection is reduced after menopause as circulating estrogen levels decline. Estrogens play a critical role in maintaining metabolic health. In women who undergo bilateral oophorectomy before age 45, hormone therapy attenuates many of the adverse metabolic changes that occur after surgery^3^. Recent meta-analyses of clinical trials show that menopausal hormone therapy (MHT), including both estrogen only and combined estrogen-progestin regimens, improves glucose homeostasis and related metabolic parameters in postmenopausal women^4^. However, regarding ASCVD events, MHT approaches have not shown consistent benefits in postmenopausal women^5–7^. Understanding the cardiovascular effects of MHT in older women has become increasingly important in light of the FDA’s recent decision to remove several boxed warnings from MHT products^8^.

Cardiovascular benefits observed in early years of MHT trials led to the “Timing Hypothesis,” which posits that MHT effects depend on the timing of initiation relative to age and years since menopause^9^. Estradiol treatment slowed the progression of carotid-intima-media thickness in women who initiated MHT within 6 years after menopause, but not in those who initiated treatment 10 or more years after menopause^10^. In postmenopausal women with established coronary atherosclerotic disease^5,6^ or pre-existing atherosclerosis^7,9,11–13^, despite favorable changes in lipid profiles, MHT did not reduce the incidence of primary cardiovascular outcomes, including myocardial infarction and coronary heart disease death, nor did it improve secondary cardiovascular outcomes, including cardiac arrest, stroke, and transient ischemic attack^5,6^. Notably, increases in inflammatory markers, such as C-reactive protein, were observed following MHT^7,11^, while inflammation has been recognized as a chronic risk factor for cardiovascular diseases.

Estrogens exert their biological effects through three receptors, estrogen receptor α, β, and the G protein-coupled estrogen receptor (ERα, ERβ, and GPER). Among these, ERα is the principal mediator of metabolic homeostasis, playing a critical role in maintaining glucose and lipid homeostasis^14,15^. We have previously demonstrated that hepatic ERα signaling was required to limit liver fat accumulation and maintain glucose homeostasis during high-fat diet feeding^16,17^. Another research group showed that estrogen treatment improved metabolism but induced pro-inflammatory effects in ovariectomized mice fed a high-fat diet^18^.

To understand the underlying mechanisms for the discordance between metabolic improvements of MHT and its limited cardiovascular benefits, we developed mouse models of pre-established atherosclerosis incorporating 17β-estradiol (E_2_) treatment following surgical menopause (ovariectomy, OVX) in young and older hyperlipidemic mice. Our findings demonstrate that E_2_ treatment following OVX in older *Ldlr^−/−^* mice with established atherosclerosis fails to improve atherosclerotic burden to levels observed in sham-operated mice during lipid normalization. We further show that E_2_ treatment improves lipid metabolism but also increases systemic inflammation. Importantly, suppression of inflammation through transplantation of *Ifnγ^−/−^* bone marrow attenuates atherosclerosis in mice receiving OVX+E_2_ treatment. Mechanistically, the persistent inflammation in the older E_2_-treated mice is associated with enhanced signaling pathways involved in fatty acid oxidation, cellular stress responses, and inflammation. In contrast to older mice, the young mice exhibit markedly attenuated oxidative stress and inflammatory responses following OVX+E_2_ treatment, accompanied by reduced atherosclerosis. These age-dependent differences may be attributable, in part, to hepatic estrogen receptor remodeling and altered inflammatory response to metabolic stress. Together, our findings provide mechanistic insights into how aging-associated alterations in estrogen signaling and inflammatory responses to E_2_ may compromise the metabolic benefits of MHT, potentially contributing to ASCVD risk in older postmenopausal women.

## Materials and Methods

### Animal Studies

The mouse colonies were housed and maintained in 12-hour light/dark cycles in temperature and humidity-controlled facilities with ad-libitum access to chow and water. *Ldlr^−/^*^−^ (Jax. Strain #002207) and *Ifnγ^-/-^* (Jax. Strain #00287) mice were originally purchased from Jax lab. All mouse experiments were approved under the Institutional Animal Care and Use Committee at Vanderbilt University Medical Center and The Ohio State University.

Atherosclerosis was established in female *Ldlr^−/−^* mice by feeding a Western diet (WD, Research Diets Inc., D12079B) for 12 weeks, followed by a switch to a chow diet to normalize circulating lipid levels. A subset of mice was sacrificed 14 days after the dietary switch, when atherosclerosis peaked, and served as the baseline control group^19^. The remaining mice underwent ovariectomy followed by E_2_ treatment (OVX+E_2_), ovariectomy alone (OVX), or sham surgery, as previously described^16,20^. E_2_ was administered through subcutaneous implantation of a 90-day sustained-release 17β estradiol pellet (0.25 mg; Innovative Research, NE-21). Following an additional 10 weeks of chow feeding, mice were sacrificed after a 5-hour fast. Serum, liver, and aorta tissues were collected and stored under appropriate conditions for subsequent analyses.

In this study, inflammation was suppressed by bone marrow transplantation using donor cells from *Ifnγ^−/−^* mice, as previously discribed^21^. Briefly, following 12 weeks of WD-feeding, mice that underwent lethal irradiation (900 rad) were reconstituted with bone marrow (*Ifnγ^−/−^*^BM^) from 6-8-week-old donor mice. Mice were allowed to recover on a chow diet for 2 weeks before undergoing surgeries for hormone treatments.

### Lipid, insulin, and circulating cytokine assessments

Fast protein liquid chromatography (FPLC) using a Superose 6 size-exclusion column (GE Healthcare) was performed to separate lipoprotein fractions, including VLDL, LDL and HDL, from serum samples pooled from 2-3 mice. Cholesterol and triglyceride concentrations in whole serum and FPLC fractions were measured using enzymatic colorimetric assays (Cholesterol Reagent and Triglycerides GPO Reagent kits, Raichem), as previously described^22^.

Fasting serum insulin concentrations were measured using a mouse insulin ELISA kit (Crystal Chem, Cat. # 90080). Serum IFNγ and IL-6 concentrations in aging and *Ifnγ^−/−BM^* recipient mice were quantified using mouse IFNγ (R&D Systems, Cat. # MIF00) and IL-6 (R&D Systems, Cat. # M600B) ELISA kits.

Liver lipid content in all mice and serum cytokine levels in young mice were measured by the Hormone & Analytical Service Core at Vanderbilt University Medical Center.

### HDL function assays

HDL function was evaluated with ApoB-free serum as we reported before^23,24^.

Antioxidant capacity was quantified using an OxiSelect Total Antioxidant Capacity Assay Kit (Cell Biolabs, Cat. # STA-360). Total antioxidant power was converted and represented as Copper Reducing Equivalents (CRE), as previously described^23,24^.

Cholesterol efflux capacity was assessed using THP-1-derived macrophages with a fluorescence-based cholesterol efflux assay kit (Abcam, ab196985), following previously described protocols^23,24^. Briefly, THP-1 cells were seeded into 96-well plates and differentiated into macrophages by treatment with phorbol 12-myristate 13-acetate (PMA, 10ng/mL, Sigma-Aldrich) for 48 hours. Following differentiation, cells were washed twice with PBS and labelled with fluorescent cholesterol using the labeling reagent supplied with the kit for 1 hour at 37°C. Cells were then washed twice with PBS and incubated with RPMI media containing apoB-deleted serum from experimental mice for 4 hours at 37°C. Culture supernatants were collected, and cells were washed and lysed using Lysis Buffer I (Abcam). Fluorescence in the supernatants and cell lysates was measured using a SpectraMax M2 microplate reader at an emission wavelength of 523nm. Cholesterol efflux capacity was calculated as the percentage of fluorescence released into the culture supernatants relative to the total fluorescence measured in supernatants and cell lysates.

### Characterization of atherosclerotic burden

Atherosclerotic burden was characterized by assessing lesion size, inflammation, and plaque vulnerability. Lesion area was quantified in frozen cross-sections of the aortic root stained with Oil Red O, as previously described^20,23,25^. Lesion inflammation was evaluated by immunofluorescence staining for CD68-positive (CD68^+^) macrophages^19,25^. Briefly, sections were fixed in cold acetone for 10 minutes, washed twice with PBS, and blocked with Background Buster (Innovex) for 1 hour at 37°C. Sections were then incubated with a biotin-conjugated anti-CD68 antibody (Calbiochem) overnight at 4°C. Following three washes with PBS, sections were incubated with FITC-conjugated streptavidin (Life Technologies) for 1 hour at 37°C, washed, and imaged. Plaque vulnerability was analyzed by quantifying necrotic core area in H&E-stained sections, as previously described^25^. Images were captured using an Olympus IX81 microscope and analyzed using the KS300 imaging system (Kontron Elektronik GmbH) or ImageJ software.

### Immunoblotting

Immunoblotting with liver tissues was performed as we reported previously^21^. The primary antibodies for pACC (3661), ACC (3676), FASN (3189), AMPKβ (4150), pAMPKβ (4186), pAMPKα (2531), and AMPKα(2603) were from Cell Signaling. The primary antibodies for PGC1a (ab54481) and HNJ2 (ab48506) were from Abcam. Liver β-actin was detected with mouse anti-human actin (MCA5775GA) from Bio-Rad. Secondary antibodies were from LI-COR biotech (Lincoln, NE). All primary antibodies were diluted at 1:1000 and incubated at 4 °C overnight. All secondary antibodies were diluted at 1:10,000 and were incubated at room temperature for 1 hour. Images were acquired using an LI-COR Odyssey infrared imaging system (LI-COR Biotechnology). Blot densities were quantified with Image J.

### Liver Bulk RNAseq and Gene Expression

Liver mRNA was assessed through bulk-RNAseq and qPCR for gene expression changes. Mice were sacrificed, and liver samples were flash-frozen in liquid nitrogen immediately and stored in -80°C until later use. Liver tissue (25mg) was bead-homogenized and total RNA was isolated according to manufacturer’s instructions (Direct-zol RNA Miniprep Kit, Zymo Research). Isolated RNA was treated with DNAse I and submitted (∼80ng/µL) to Vanderbilt Technologies for Advanced Genomics (VANTAGE) core laboratory for Next-Generation sequencing. RNA samples were sequenced using multiplexed Paired-End 150bp on the Illumina NovaSeq 6000. Quality control (QC) was evaluated at different levels including RNA quality, raw read data, alignment, and gene expression. Raw paired-end reads were mapped to the mouse reference genome mm10 using STAR 2.7.3^26^. Feature count was used to calculate raw read counts and downstream analysis^27^. Differential gene expression and functional enrichment analyses were performed in R (version 3.6) with several Bioconductor packages: DESeq2, clusterProfiler. Comparisons were made between the sham and OVX+E_2_-treatment groups. Differentially expressed genes were defined as genes with a Benjamini–Hochberg-adjusted *P* value (FDR) < 0.05 and an absolute fold change >1.5 (|log2 fold change| >0.585). Results for specific comparison were extracted by combinations of coefficients with DESeq2. A 5% false discovery rate threshold was used for significant results filtering. Gene expression was validated with qPCR, which was performed in duplicate using TaqMan primers (Supplemental Table 1) and reagents from Life Technologies.

### Statistical Analysis

Data are presented as mean SEM, unless otherwise indicated. Statistical differences among groups were assessed using ANOVA with appropriate post hoc multiple-comparison tests, as specified in the corresponding figure legends. A *P* value <0.05 was considered statistically significant. Statistical methods used for RNA-seq analysis are described in the preceding section.

## Results

### 17β-estradiol (E_2_) treatment fails to inhibit atherosclerosis following ovariectomy (OVX) in older *Ldlr*^−/−^ mice

To model MHT in the setting of established atherosclerosis, atherosclerosis was established in 11-month-old female *Ldlr^−/−^* mice by 3 months of Western diet (WD)-feeding (Fig. 1A). Mice were then switched to a chow diet to minimize lipid-related risk. Two weeks after the diet switch, when atherosclerotic burden peaked^20^, one group of mice was euthanized for baseline plaque evaluation. The remaining mice were divided into three groups matched for body composition: sham surgery, OVX, or E_2_ treatment following OVX (OVX+E_2_). After 10 weeks of hormone treatments while maintained on a chow diet, mice were euthanized for metabolic and atherosclerotic assessments. Compared with WD-feeding, chow diet-feeding reduced fasting cholesterol and triglyceride (TG) concentrations by ∼80% and ∼37% (*P*<0.001), respectively, accompanied by improved lipoprotein profiles across all hormone treatment groups (Fig. 1, B-E and Suppl. Fig. 1, A-F). Ovariectomy increased fasting TG concentrations in OVX mice relative to sham mice; E_2_ treatment reduced fasting TG concentrations following OVX (Fig. 1C). Chow diet-feeding decreased atherogenic VLDL-cholesterol by ∼98% (418.25 vs. 9.28 mg/dL, *P*<0.0001) and LDL-cholesterol by ∼72% (893.8 vs. 167.2 mg/dL, *P*<0.0001) compared with WD-feeding conditions (Suppl. Fig. 1, A-B). Similarly, chow diet-feeding decreased VLDL-TG by ∼88% (91.3 vs 10.8 mg/dL, *P*<0.0001) and LDL-TG by ∼ 75% (135.1 vs 34.3 mg/dL, *P*<0.0001) compared with WD-feeding conditions (Suppl. Fig. 1, D-E). Chow diet-feeding also reduced fasting circulating insulin at baseline and did not further reduce insulin levels among hormone treatment groups (Fig. 1F). These findings demonstrate that atherogenic hyperlipidemia and insulin homeostasis were substantially improved after switching from WD to a chow diet and suggest that E_2_ treatment improved lipid metabolism in ovariectomized *Ldlr^−/−^*mice.

**Figure 1.**
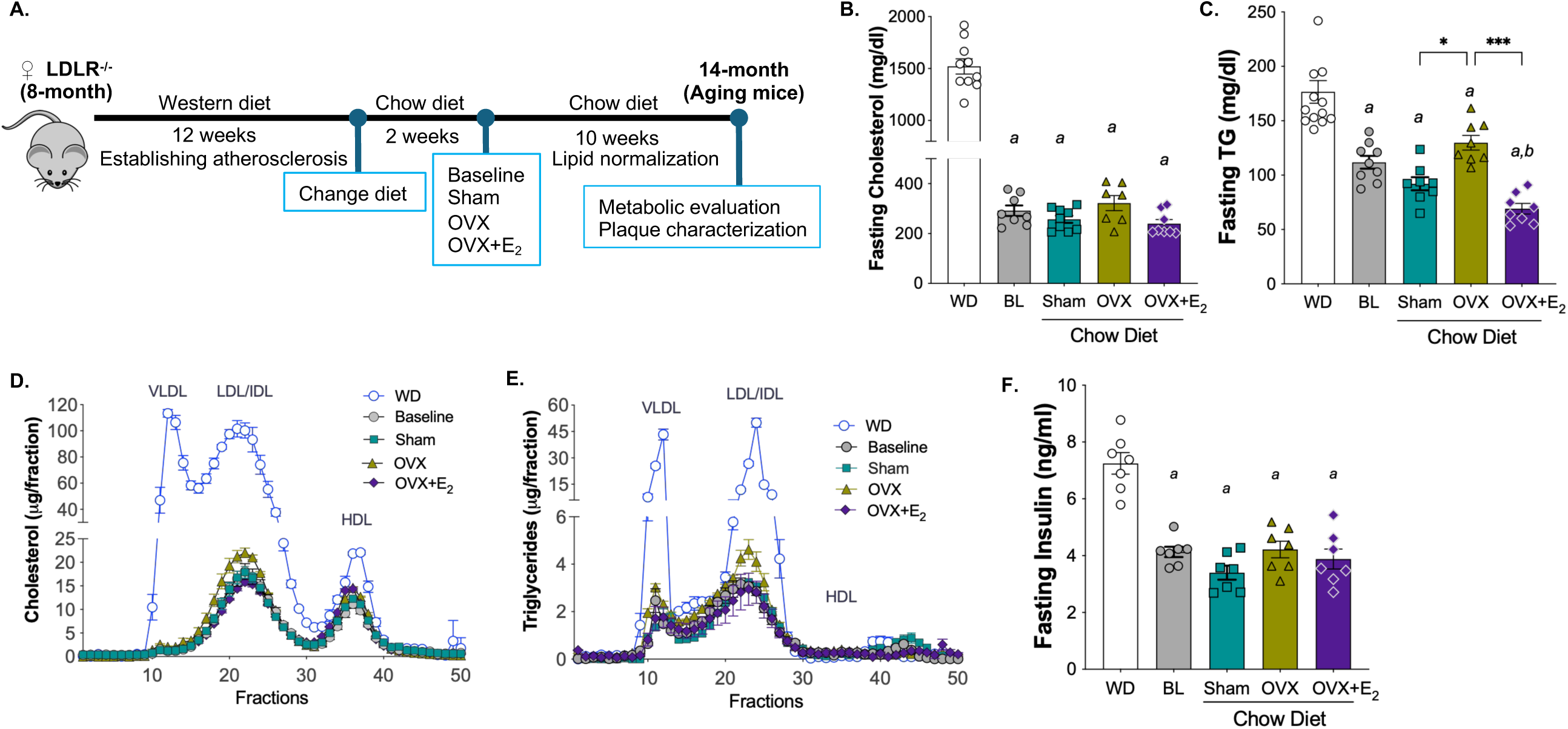
E_2_ treatment following OVX improves circulating lipid profiles and reduces insulin concentrations during lipid management in aging *Ldlr^−/−^* mice. **A**. Schematic of the study design. **B.** Fasting serum total cholesterol levels. **C.** Fasting total triglyceride (TG) levels. **D-E**. Lipoprotein cholesterol (D) and TG (E) contents determined by fast protein liquid chromatography (FPLC). **F.** Fasting serum insulin concentrations. Data are presented as mean ± SEM (n >=7/group). Statistical analyses were performed using one-way ANOVA with Tukey’s multiple comparisons test. *a*, *P*<0.05 compared with Western diet (WD) conditions; *b*, *P*<0.05 compared with baseline (BL). *, *P* < 0.05; **, *P* < 0.01; ***, *P* < 0.001.

To determine the impact of E_2_ treatment on atherosclerotic burden, we analyzed aortic root lesions using Oil Red O staining to assess lesion size, CD68^+^ immunofluorescence to assess lesional inflammation, and plaque vulnerability. Overall lesion size in hormone treatment groups did not differ significantly from baseline. However, lesion area was enlarged by OVX compared with sham operation, whereas E_2_ treatment following OVX did not reduce lesion size to the level observed in sham mice (Fig. 2, A-B). CD68^+^ area, a marker of macrophage accumulation^20,25^, was reduced by ∼59% in sham mice compared with baseline and remained elevated in OVX and OVX+E_2_ mice (Fig. 2, C-D). Plaque vulnerability, assessed by necrotic core area, showed a similar pattern to CD68^+^ staining. Necrotic area was reduced only in sham mice compared with baseline and remained elevated in both OVX and OVX+E_2_ mice (Fig. 2, E-F). These data indicate that estradiol treatment did not improve atherosclerotic burden in 11-14 months old ovariectomized *Ldlr^−/−^* mice to the level exhibited in sham-operated mice. Hereafter, the hyperlipidemic *Ldlr^−/−^* mice that underwent 3 months of WD-feeding are referred to as the aging cohort to distinguish them from the other two study cohorts.

**Figure 2.**
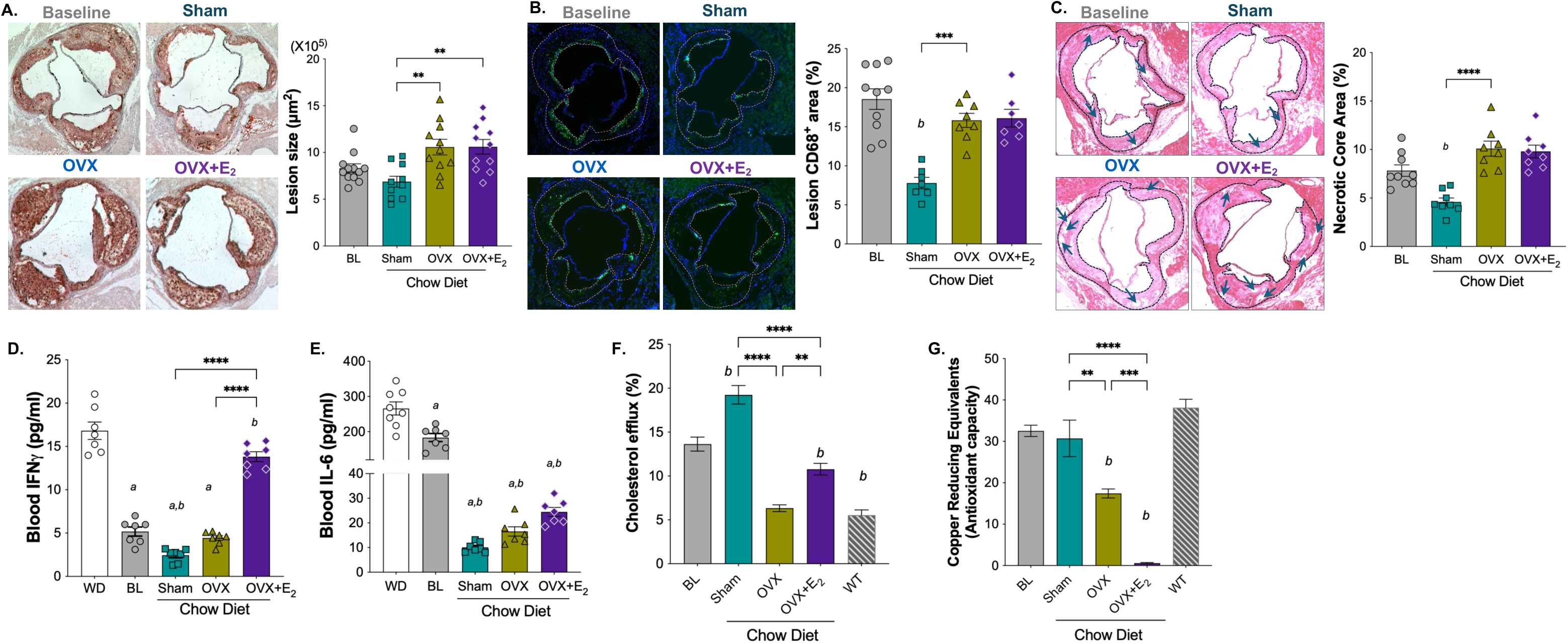
E_2_ treatment following OVX fails to reduce atherosclerotic burden during lipid management in aging *Ldlr^−/−^* mice. **A-B.** Representative Oil Red O staining of aortic root sections (A) and quantification of Oil Red O-positive area (B). **C.** Representative images of CD68^+^ immunofluorescent staining, CD68^+^ staining is shown in green, and nuclei are counterstained with DAPI (blue**). D.** Quantification of CD68^+^ area within the lesion. **E.** Representative images of H&E-stained aortic root sections. Lesion areas are outlined with dark blue dashed lines, and representative necrotic cores are indicated by arrows**. F.** Quantification of necrotic core areas. **G-H.** Circulating concentrations of the inflammatory cytokines IFNγ (G) and IL-6 (H). **I-J.** HDL cholesterol efflux capacity (I) and antioxidant capacity (J) were assessed as described in the Methods. HDL particles isolated from 3-month-old C57BL/6 wild type (WT) female mice fed a chow diet were used as controls. Data are presented as mean ± SEM (n>=7/group). Statistical analyses were performed using one-way ANOVA with Tukey’s multiple comparisons test. *a,* P<0.05 compared to Western Diet (WD) feeding conditions; *b*, *P*<0.05 compared with baseline (BL) conditions; *, *P*<0.05; **, *P*<0.01; ***, *P*<0.001; ****, *P*<0.0001.

We next evaluated systemic inflammation by measuring circulating cytokine levels, given the association between inflammation and cardiovascular events in MHT trials^7,11^. Chow diet-feeding reduced circulating IFNγ concentrations at baseline, with a further reduction in sham-operated mice but not in OVX mice (Fig. 2G). Circulating IFNγ concentrations remained elevated in OVX+E_2_ mice, reaching approximately fourfold the levels observed in sham mice (Fig. 2G). Circulating IL-6 levels were reduced at baseline following the diet switch and declined further in all hormone treatment groups (Fig. 2H). Although IL-6 levels were higher in OVX+E_2_ mice than in OVX and sham mice, these differences did not reach statistical significance (Fig. 2H).

Because increased inflammation can compromise HDL function^24,28^, we next assessed the major anti-atherogenic functions of HDL. HDL particles facilitate the removal of excess cholesterol from the subendothelial space in the artery^23^. HDL cholesterol efflux capacity was improved following lipid normalization in sham mice compared with baseline; however, this improvement was not observed in either OVX or OVX+E_2_ mice (Fig. 2I). HDL particles also exert antioxidant function that limits LDL oxidation, and LDL oxidation is an important atherogenic process during hyperlipidemia^28^. HDL antioxidant capacity was unchanged in sham mice but was reduced by ∼64% in OVX mice compared with baseline (Fig. 2J). Notably, HDL antioxidant capacity was almost completely abolished in OVX+E_2_ mice (Fig. 2J). Thus, associated with persistently elevated circulating IFNγ levels, the anti-atherogenic functions of HDL particles were markedly impaired in OVX+E_2_ mice compared with sham mice. These data demonstrate that, despite favorable changes in the lipid profile, menopausal E_2_ treatment fails to reduce established atherosclerosis in aging *Ldlr^−/−^* mice following lipid normalization. Instead, E_2_ treatment is associated with persistent systemic and plaque inflammation, a more vulnerable plaque phenotype, and impaired anti-atherogenic HDL functions.

### Targeting inflammation restores the ability of E_2_ treatment to reduce established atherosclerosis in aging ovariectomized *Ldlr^−/−^* mice

To determine whether persistent inflammation contributes to a lack of cardiovascular benefit of menopausal E_2_ treatment, we designed a second study to suppress inflammation, particularly that arising from hematopoietic cells and recruited to atherosclerotic lesions^29,30^. Similar to the first study, atherosclerosis was established in 11-month-old female *Ldlr^−/−^* mice by 3 months of WD feeding. In contrast to the first study, mice were subjected to lethal irradiation and reconstituted with *Ifn*γ^−/−^ bone marrow cells (*Ifnγ*^−/−BM^) at the time of the switch from WD to a chow diet (Fig. 3A). Two weeks after the diet switch, a group of mice that did not undergo bone marrow transplantation was euthanized as the baseline control, while the bone marrow recipient mice underwent hormone treatment procedures and were maintained on a chow diet for an additional 10 weeks (Fig. 3A).

**Figure 3.**
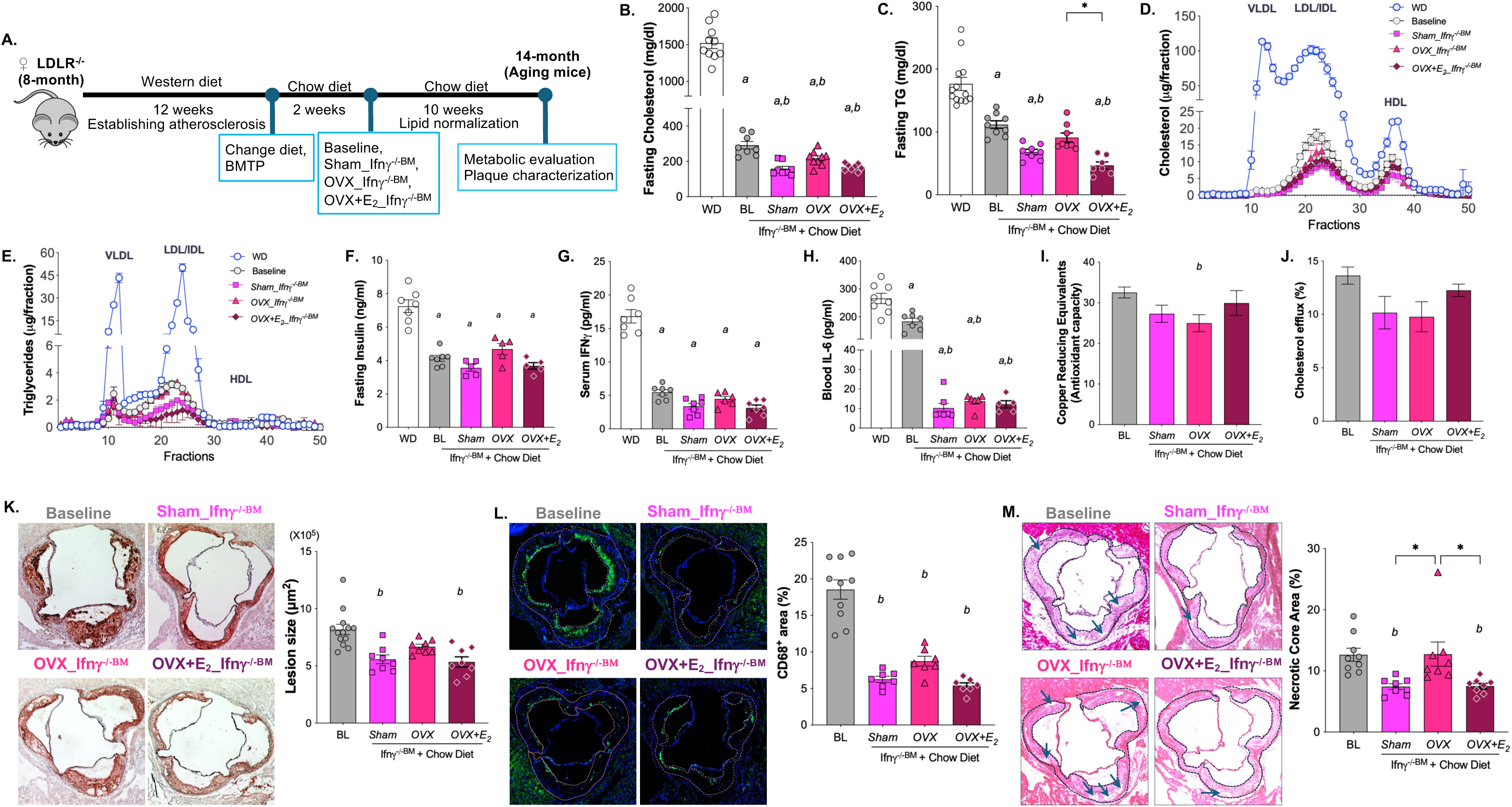
Suppression of IFNγ-mediated inflammation in myeloid cells promotes atherosclerosis regression in aging *Ldlr^−/−^* mice treated with E_2_ following OVX during lipid management. **A.** Experimental design for suppression of inflammation using *Ifnγ^−/−^* bone marrow transplantation (*Ifnγ*^−/−BM^). **B-C.** Fasting serum cholesterol (B) and TG (C) concentrations. **D-E.** Lipoprotein cholesterol (D) and TG (E) contents determined by FPLC. **F-I.** Circulating concentrations of insulin (F), IFNγ (G), and IL-6 (H). **I-J.** HDL antioxidant capacity assessed by copper reducing equivalents (I) and HDL cholesterol efflux capacity (J). **K.** Representative Oil Red O staining of aortic root sections (left) and quantification of Oil Red O-positive area (right). **L.** Left: Representative images of CD68^+^ immunofluorescent staining, CD68^+^ staining is shown in green, and nuclei are counterstained with DAPI (blue). Right: Quantification of CD68^+^ area within the lesion. **M.** Left: Representative images of H&E-stained aortic root sections. Lesion areas are outlined with dark blue dashed lines, and representative necrotic cores are indicated by arrows. Right: Quantification of necrotic core areas. Data are presented as mean ± SEM (n>=7/group). Statistical analyses were performed using one-way ANOVA with Tukey’s multiple comparisons test. *a*, *P*<0.05 compared to Western Diet (WD) feeding conditions; *b*, *P*<0.05 compared with baseline (BL) conditions; *, *P*<0.05.

Metabolic measurements obtained during WD-feeding were pooled across the 1^st^ and 2^nd^ cohort studies, as were data from the baseline groups of both cohorts. Consistent with the lipid-lowering effects of chow diet-feeding observed in the 1^st^ study (aging cohort), fasting serum cholesterol and TG concentrations, lipid contents across lipoprotein fractions, and circulating insulin concentrations were reduced at baseline compared with WD conditions (Fig. 3, B-F). Circulating insulin concentrations remained similar to baseline levels across hormone treatment groups, whereas cholesterol and TG levels were further reduced in hormone treatment groups, except for TG levels in OVX mice (Fig. 3, B-F). In contrast to observations in the aging cohort, LDL-cholesterol and LDL-TG levels were further reduced across hormone treatment groups compared with baseline (Fig. 3, D–E, Suppl. Fig. 1, H and K). These additional lipid-lowing effects may reflect, at least in part, the contribution of LDLR-expressing hematopoietic cells derived from *Ifn*γ^−/−^ donor bone marrow following transplantation, thereby enhancing clearance of circulating lipids.

Changes in circulating IL-6 levels were similar to those observed in the aging cohort. In contrast, circulating IFNγ levels were reduced in OVX+E_2_ mice following *Ifnγ*^−/−BM^ transplantation and were accompanied by improved HDL functions for cholesterol efflux and antioxidant capacity (Fig. 3, F-J). Importantly, atherosclerotic lesion size, macrophage content (CD68^+^ area), and necrotic core area were also reduced in OVX+E_2_ mice to levels comparable to those observed in sham mice (Fig. 3, K-M). Together, these findings demonstrate that suppression of hematopoietic IFNγ-mediated inflammation restores HDL function and the cardiovascular benefit of E_2_ treatment in aging ovariectomized *Ldlr^−/−^* mice during lipid normalization. These results further support that persistent inflammation could be a major barrier to the cardiovascular protective effects of menopausal E_2_ treatment in the setting of established atherosclerosis.

### E_2_ treatment following OVX reduces established atherosclerosis in young adult *Ldlr^−/−^* mice

We previously reported the metabolic benefits of E_2_ treatment following OVX in young adult female mice (3-9 months old) during high-fat diet feeding^16^. To determine whether age influences the cardiovascular response to estradiol treatment, we next performed a study in young adult mice. In the young mouse model, atherosclerosis was established in 6-month-old female *Ldlr^−/−^* mice (Fig. 4A). Following hormone treatments and lipid normalization, mice were euthanized at 9 months old, when reproductive capacity remains preserved.

**Figure 4.**
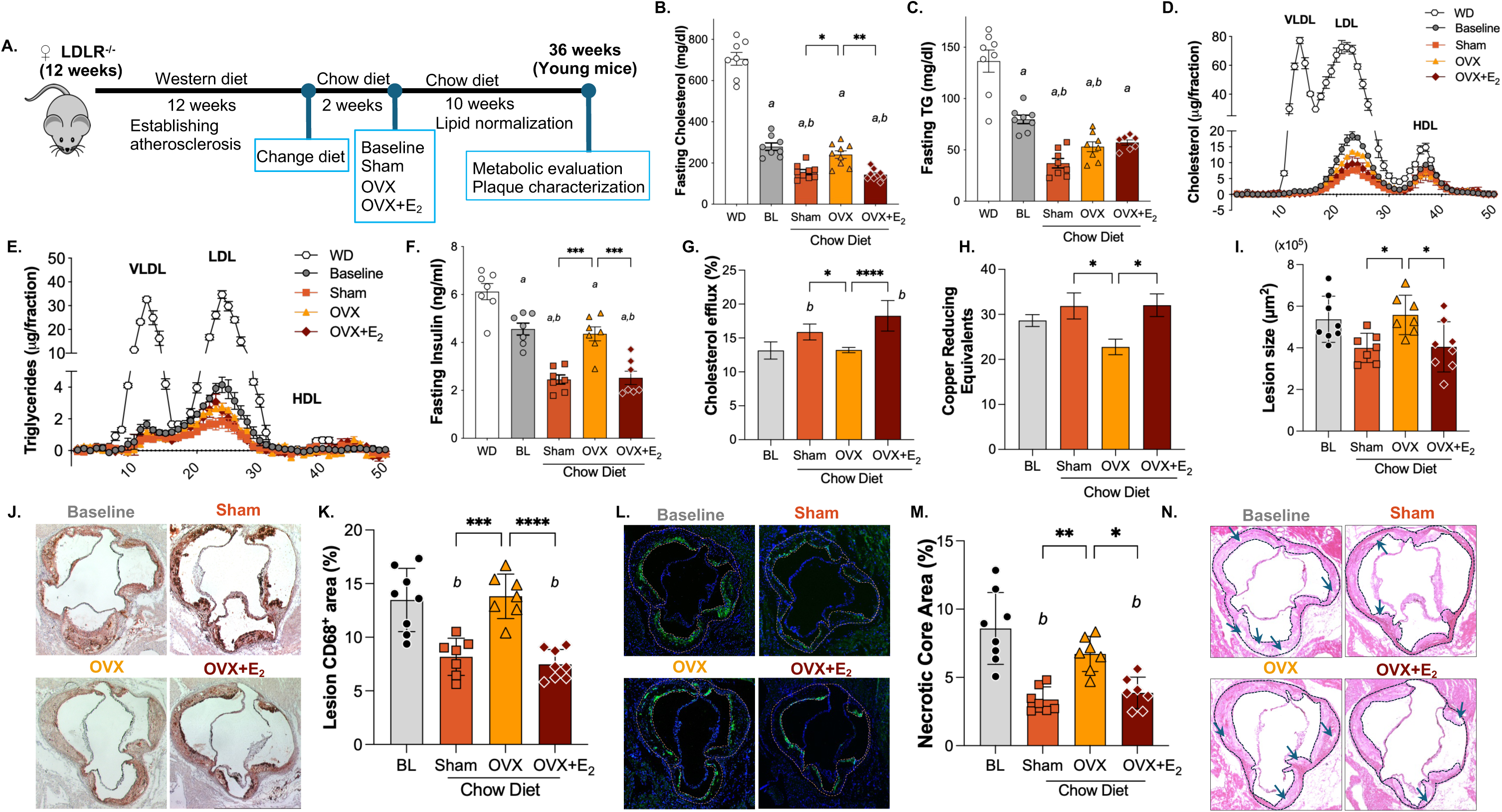
E_2_ treatment following OVX reduces atherosclerotic burden and improves HDL function in young *Ldlr^−/−^* mice during lipid management. **A.** Hormone therapy was modeled in younger adult female *Ldlr^−/−^* mice that maintained reproductive function throughout the study. **B-C.** Fasting serum cholesterol (B) and triglyceride (C) concentrations were similarly reduced in sham and OVX+E_2_ mice. **D-E.** Lipoprotein distribution of cholesterol (D) and triglyceride (E) determined by FPLC. **F.** Fasting insulin concentrations were similarly reduced in sham and OVX+E_2_ mice. **G.** HDL cholesterol efflux capacity. **H.** HDL antioxidant capacity assessed by copper reducing equivalents. **I-J.** Aortic root lesion area was quantified by Oil Red O staining (I), with representative images from each group shown in J. **K-L.** CD68^+^ area within atherosclerotic lesions was reduced in sham and OVX+E_2_ mice (K), with representative images shown in L. **M-N**. Necrotic core area was reduced in sham and OVX+E_2_ mice (M), with representative images shown in N. Data are presented as mean ± SEM (n>=7/group). Statistical analyses were performed using one-way ANOVA with Tukey’s multiple comparisons test. *a*, *P*<0.05 compared to Western Diet (WD) feeding conditions; *b*, *P*<0.05 compared with baseline (BL) conditions; *, *P*<0.05; **, *P*<0.01; ***, *P*<0.001; ****, *P*<0.0001.

Similar to observations in older mice, the transition from WD to chow diet substantially improved lipid profiles. Fasting cholesterol and TG concentrations were reduced by ∼61% and ∼42%, respectively, at baseline compared with WD conditions (Fig. 4B-C). Fasting cholesterol levels were further reduced in sham and OVX+E_2_ mice, whereas fasting TG concentrations were further reduced in sham and OVX mice, compared with baseline (Fig. 4, B-C). VLDL-and LDL-associated cholesterol were reduced by ∼98% and ∼77%, respectively, at baseline and were reduced by an additional ∼50% in sham and OVX+E_2_ mice compared with baseline (Fig. 4D, Suppl. Fig. 1, M-O). Similar improvement in VLDL- and LDL-associated TG presented at baseline as well as in sham and OVX+E_2_ mice (Fig. 4E, Suppl. Fig. 1, P-R). Fasting insulin levels were lower at baseline than under WD conditions and were further reduced in sham and OVX+E_2_ mice, but not in OVX mice (Fig. 4F).

In contrast to the aging cohort (Fig. 2, G-H), young mice exhibited reduced circulating IFNγ and other inflammatory marker concentrations following menopausal E_2_ treatments (Table 1). This reduction in systemic inflammation was accompanied by improved HDL antioxidant and cholesterol efflux functions (Fig. 4, G-H). Overall atherosclerotic lesion area across the hormone treatment groups did not differ significantly from baseline; however, lesion area was smaller in both sham and OVX+E_2_ mice than in OVX mice (Fig. 4, I-J). More pronounced reductions were observed in plaque inflammation and vulnerability in both sham and OVX+E_2_ mice compared with baseline and OVX mice (Fig. 4, K-N). These data indicate that sham-operation and OVX+E_2_ treatment similarly improved atherosclerotic plaque characteristics in young mice.

**Table 1.**
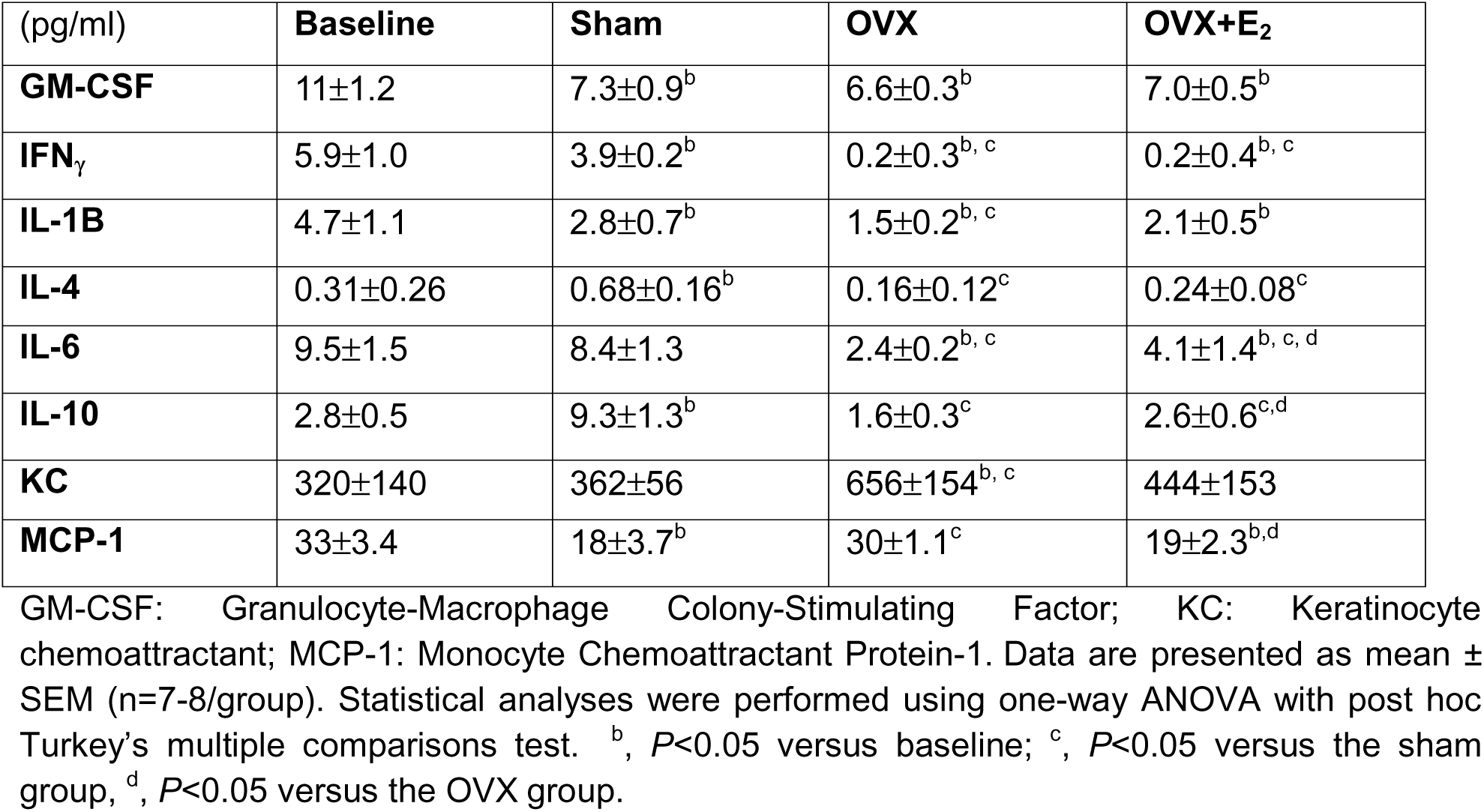
Serum cytokine concentrations from young mouse groups.

| (pg/ml) | **Baseline** | **Sham** | **OVX** | **OVX+E<sub>2</sub>** |
| --- | --- | --- | --- | --- |
| **GM-CSF** | 11±1.2 | 7.3±0.9<sup>b</sup> | 6.6±0.3<sup>b</sup> | 7.0±0.5<sup>b</sup> |
| **IFN<sub>γ</sub>** | 5.9±1.0 | 3.9±0.2<sup>b</sup> | 0.2±0.3<sup>b, c</sup> | 0.2±0.4<sup>b, c</sup> |
| **IL-1B** | 4.7±1.1 | 2.8±0.7<sup>b</sup> | 1.5±0.2<sup>b, c</sup> | 2.1±0.5<sup>b</sup> |
| **IL-4** | 0.31±0.26 | 0.68±0.16<sup>b</sup> | 0.16±0.12<sup>c</sup> | 0.24±0.08<sup>c</sup> |
| **IL-6** | 9.5±1.5 | 8.4±1.3 | 2.4±0.2<sup>b, c</sup> | 4.1±1.4<sup>b, c, d</sup> |
| **IL-10** | 2.8±0.5 | 9.3±1.3<sup>b</sup> | 1.6±0.3<sup>c</sup> | 2.6±0.6<sup>c, d</sup> |
| **KC** | 320±140 | 362±56 | 656±154<sup>b, c</sup> | 444±153 |
| **MCP-1** | 33±3.4 | 18±3.7<sup>b</sup> | 30±1.1<sup>c</sup> | 19±2.3<sup>b, d</sup> |

GM-CSF: Granulocyte-Macrophage Colony-Stimulating Factor; KC: Keratinocyte chemoattractant; MCP-1: Monocyte Chemoattractant Protein-1. Data are presented as mean ± SEM (n=7-8/group). Statistical analyses were performed using one-way ANOVA with post hoc Turkey's multiple comparisons test. <sup>b</sup>, *P*<0.05 versus baseline; <sup>c</sup>, *P*<0.05 versus the sham group, <sup>d</sup>, *P*<0.05 versus the OVX group.

Collectively, findings across aging and young cohorts demonstrate that menopausal estradiol treatment exerts beneficial metabolic and cardiovascular effects in young *Ldlr^−/−^* mice following lipid normalization. In contrast to aging mice, young mice exhibited reduced systemic and plaque inflammation, improved HDL function, and reduced atherosclerotic burden during E_2_ treatment following OVX. These findings indicate that the cardiovascular protective effects of estradiol treatment are preserved in young mice but become compromised with aging.

### E_2_ treatment following OVX promotes hepatic fatty acid metabolism favoring oxidation in both aging and young mice

We previously reported that HDL function correlates with hepatic lipid metabolism^24^ and that estradiol treatment following OVX promotes hepatic fatty acid oxidation in young adult mice during high-fat diet feeding, in part through increased phosphorylation of acetyl-CoA carboxylase (ACC; pACC)^16,17^. To better understand the mechanisms underlying persistent inflammation and impaired HDL function associated with OVX+E_2_ treatment in aging mice, we assessed hepatic lipid content and proteins involved in lipid metabolic pathways. Chow diet-feeding reduced hepatic cholesterol content in all hormone treatment groups compared with baseline and reduced hepatic TG content in sham and OVX+E_2_ mice, in both aging and young cohorts (Fig. 5, A-D). In the aging cohort, pACC protein levels were higher in OVX+E_2_ mice, whereas total ACC protein levels were higher in OVX mice (Fig. 5, E-G). Fatty acid synthase (FASN) protein levels did not differ significantly among groups (Fig. 5, E and H). The increase in total ACC in OVX mice was consistent with a metabolic state favoring lipogenesis and fat accumulation following ovariectomy. Interestingly, protein levels of peroxisome proliferator-activated receptor γ coactivator 1-α (PGC1α), a master regulator of hepatic oxidative metabolism, were increased following lipid normalization in all hormone treatment groups compared with baseline and were higher in OVX+E_2_ than in sham mice (Fig. 5, E and I). In the young cohort, chow diet-feeding reduced pACC protein levels in all three hormone treatment groups; however, pACC levels remained higher in OVX+E_2_ mice than in sham and OVX mice, whereas total ACC protein levels remained elevated in OVX mice (Fig 5, J-L). Furthermore, changes in FASN protein levels paralleled those of total ACC, with higher FASN levels in OVX mice than in sham and OVX+E_2_ mice (Fig. 5, J and M). These findings suggest that the increased hepatic TG content in young OVX mice is likely contributable to enhanced lipogenic signaling through ACC and FASN. PGC1α protein levels were reduced in sham mice compared with baseline but remained elevated in OVX and OVX+E_2_ mice (Fig. 5, J and N). Thus, E_2_ treatment following OVX maintained molecular pathways favoring fatty acid oxidation in both aging and young cohorts.

**Figure 5.**
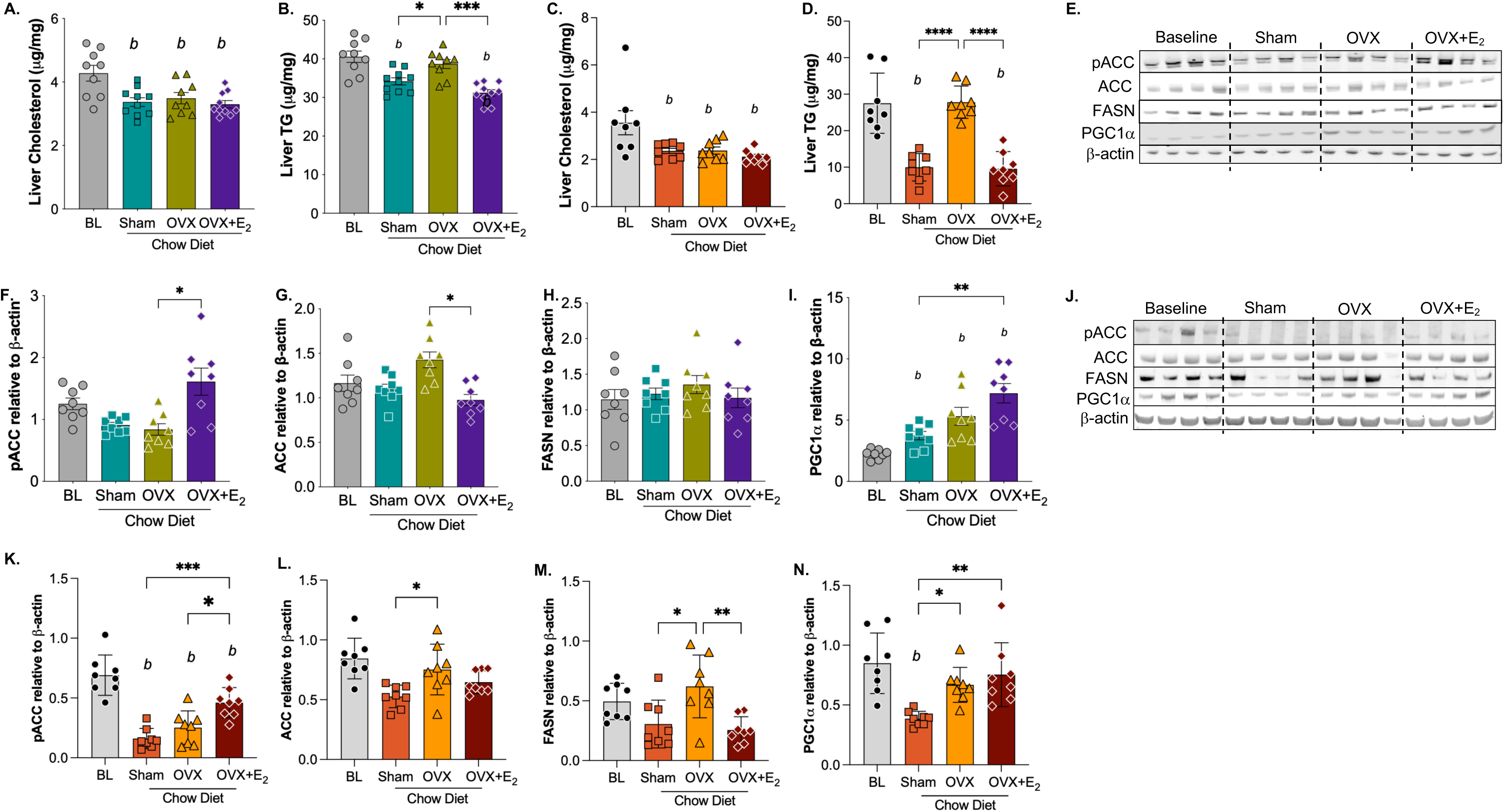
E_2_-medicated regulation of liver TG content and ACC phosphorylation is preserved with aging. **A.** Liver cholesterol content in the aging cohort. **B.** Liver TG content in the aging cohort. **C.** Liver cholesterol content in the young cohort. **D.** Liver TG content in the young cohort. Immunoblot analysis and quantification of liver phosphorylated ACC (pACC), total ACC, FASN and PGC1α with immunoblotting in the aging cohort shown in **E-I** in the young cohort shown in **J-N.** Data are presented as mean ± SEM (n>=7/group). Statistical analyses were performed using one-way ANOVA with Tukey’s multiple comparisons test. *b*, *P*<0.05 compared with baseline (BL) conditions; *, *P*<0.05; **, *P*<0.01; ***, *P*<0.001; ****, *P*<0.0001.

ACC phosphorylation is regulated by the heterotrimeric AMP-activated protein kinase (AMPK) complex^31^, which consists of catalytic α and regulatory β and γ subunits^32,33^. Phosphorylation of AMPKα increases its catalytic activity toward downstream substrates, including ACC, whereas the β subunit primarily serves as a scaffold for complex assembly and contributes to carbohydrate and glycogen sensing^32,33^. In the aging cohort, neither total AMPKα nor pAMPKα protein levels were significantly altered in hormone treatment groups compared with baseline; however, both pAMPKα and total AMPKα protein levels were higher in OVX+E_2_ mice than in sham and OVX mice (Suppl. Fig. 3, A-C). No significant differences in either pAMPKβ or total AMPKβ were observed among groups in the aging cohort (Suppl. Fig. 3, A, D-E). In contrast to the aging cohort, total AMPKα and its phosphorylation were reduced by ∼45%-68% following an additional 10 weeks of chow in the young cohort, whereas relatively higher levels of AMPKα and pAMPKα were preserved in OVX+E_2_ mice (Suppl. Fig. 3, F-H). pAMPKβ protein levels were reduced in sham and OVX mice, but not in OVX+E_2_ mice, whereas total AMPKβ protein levels did not differ significantly among groups (Suppl. Fig. 3, F, I-J). Collectively, these findings indicate that the ability of E_2_ to activate the AMPK–ACC pathway and promote a hepatic metabolic state favoring fatty acid oxidation is preserved in both aging and young cohorts. In parallel, OVX was associated with increased expression of lipogenic proteins, including ACC and/or FASN, in both cohorts, potentially contributing to hepatic lipid accumulation and increased lipotoxicity.

When inflammation was suppressed by *Ifnγ^−/−^*^BM^ transplantation, changes in hepatic lipid content and lipid metabolic proteins, including pACC, ACC, FASN, and PGC1α, across hormone treatment groups were attenuated (Suppl. Fig. 2, A-G). The increases in pAMPKα and total AMPKα observed in OVX+E_2_ mice were also attenuated following *Ifnγ^−/−^*^BM^ transplantation (Suppl. Fig. 3, K-O). These findings indicate that suppressing hepatic inflammation by eliminating IFN*γ* in bone marrow derived cells attenuated the hepatic metabolic adaptations associated with both fatty acid synthesis and oxidation during hormone treatment. This observation is consistent with our previous findings demonstrating reciprocal regulation between fatty acid oxidation and innate inflammatory signaling^21^, suggesting that suppression of inflammation through *Ifn*γ^−/−BM^ transplantation not only reduces systemic inflammation but also modifies metabolic pathways and hepatic lipid contents (Suppl. Fig. 2, A-B).

### Age-dependent hepatic estrogen receptor remodeling is associated with dysregulation of lipid peroxidation during menopausal E_2_ treatment

In mice, circulating estrogen levels do not decline with aging as markedly as they do in humans^34^, whereas estrogen receptor (ER) expression, particularly ERα, declines with aging in multiple tissues^35,36^. We therefore quantified hepatic expression of ERs in the aging and young cohorts. In the aging cohort, *Esr1* (ERα) remained the predominantly expressed estrogen receptor compared with *Esr2* (ERβ) and *Gper1* (GPER); however, *Esr1* mRNA levels were significantly lower in aging mice than in young mice (Fig. 6, A-B). In contrast to *Ers1*, *Gper1* expression was markedly higher in aging mice, whereas its expression remained low in young mice regardless of hormone treatment (Fig. 6, A and C). Both *Esr1* and *Gper1* expression responded to estrogen status in aging mice, with higher mRNA levels in sham and OVX+E_2_ mice than in OVX mice (Fig. 6, A-C). *Esr2* mRNA was detectable only at low levels and showed minimal regulation by hormone treatment in the aging cohort, whereas it was undetectable by qPCR in the young cohort (Fig. 6A). These findings demonstrate age-dependent remodeling of ER expression, characterized by reduced *Esr1* expression and relative enrichment of *Gper1* expression with aging.

**Figure 6.**
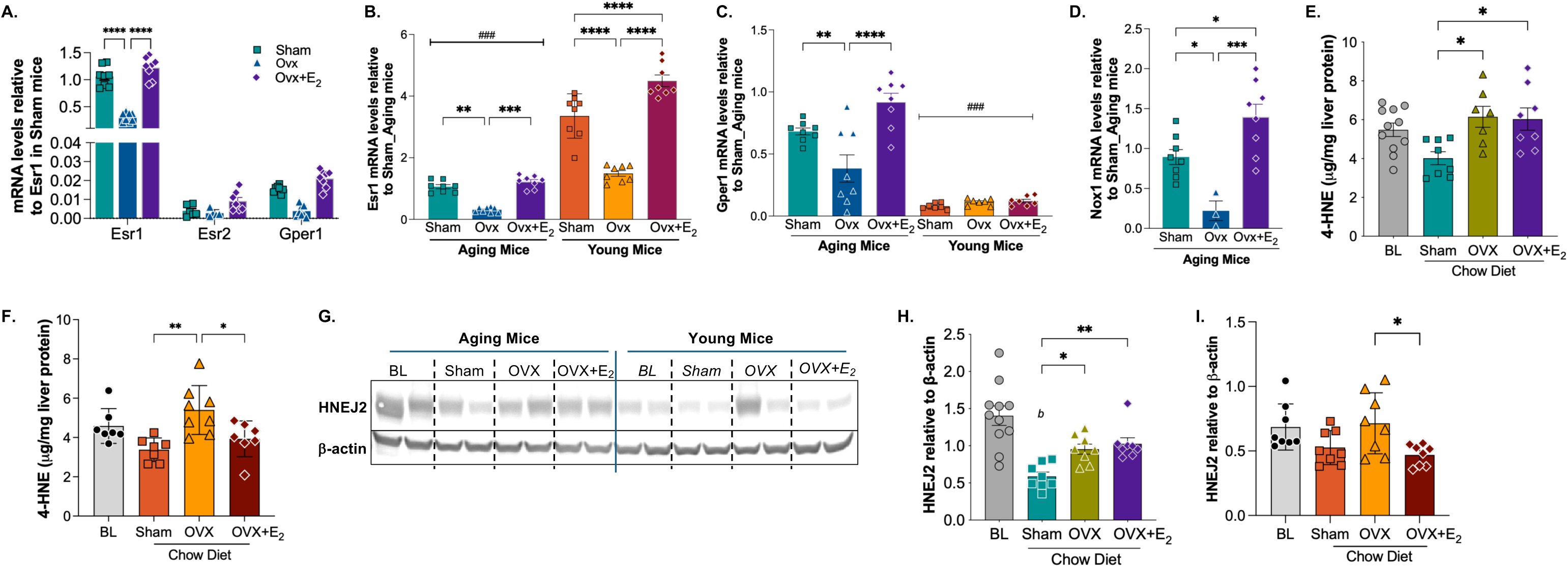
Estrogen receptor remodeling with aging is associated with increased hepatic 4-HNE and 4-HNE-protein adducts in OVX+E_2_ mice. **A.** Hepatic mRNA expression of *Esr1, Esr2,* and *Gper1* in aging mice, normalized to *Esr1* expression in sham mice. **B.** Hepatic *Esr1* expression is reduced with aging, whereas its responsiveness to estrogens is preserved in both aging and young cohorts. **C.** Hepatic *Gper1* expression increases with aging and remains estrogen responsive. D. *Nox1* expression was detectable by qPCR in the aging cohort, showing its expression is also estrogen responsive. **E-F.** Liver 4-HNE content in the aging (E) and young (F) cohorts. **G.** Representative immunoblots of liver 4-NHE-protein adducts detected using the HNEJ2 antibody in the aging and young cohorts. **H-I.** Quantification of liver HNEJ2 in the aging (E) and young (I) cohorts. Gene expression was normalized to 18s rRNA. For panels B and C, data are presented as fold change relative to the corresponding sham group in the aging cohort. Data are shown as mean ± SEM (n=7-8/group). Statistical significance was determined by two-way ANOVA with Sidak’s post hoc multiple-comparison tests for panels A-C and determined by one-way ANOVA with Tukey’s multiple comparisons test for panels D-F and H-I. ^###^, *P*<0.001 between aging and young cohorts; *b*, *P*<0.05 compared with baseline (BL) conditions; *, *P*<0.05. **, P<0.01, ***, *P*<0.001, ****, *P*<0.0001.

Previous studies demonstrated that estrogens promote NADPH oxidase 1 (NOX1)-mediated oxidative stress through constitutive activation of GPER signaling in endothelial cells^37,38^. We therefore examined hepatic *Nox1* expression in both aging and young mouse cohorts. In the aging cohort, liver *Nox1* mRNA levels were higher in sham than in OVX mice and were further increased by menopausal E_2_ treatment (Fig. 6D). In contrast, liver *Nox1* mRNA was below the limit of reliable detection by qPCR in the young cohort. NOX1-mediated reactive oxygen species (ROS) production promotes lipid peroxidation, leading to the generation of 4-hydroxynonenal (4-HNE)^37,39^. Since ROS cannot be reliably quantified in archived frozen tissues, we measured liver 4-HNE content as a marker of lipid peroxidation. In the aging cohort, overall liver 4-HNE across all hormone treatment groups did not differ from baseline (Fig. 6E). However, 4-HNE levels were elevated following ovariectomy compared with sham-operated mice and remained elevated despite menopausal E_2_ treatment (Fig. 6E). Similar to the aging cohort, liver 4-HNE content was not significantly reduced in hormone treatment groups compared with baseline in the young cohort (Fig. 6F). In contrast to the aging cohort, liver 4-HNE content was reduced during menopausal E_2_ treatment in the young cohort (Fig. 6F). Furthermore, levels of 4-HNE protein adduct detected by HNEJ2 were generally higher in the aging cohort than in the young cohort, potentially reflecting the higher hepatic TG and 4-HNE contents (Fig. 5, B and D; Fig. 6, E-I). In the aging cohort, HNEJ2 levels were increased by OVX compared with sham operation, whereas menopausal E_2_ treatment failed to reduce HNEJ2 levels. In contrast, in the young cohort, E_2_ treatment following OVX reduced HNEJ2 levels to those presented in sham mice (Fig. 6, G-I). In *Ifnγ*^−/−BM^ recipient mice, neither chow diet-feeding nor hormone treatments significantly altered liver 4-HNE content, whereas HNEJ2 levels were not reduced in sham mice but reduced in OVX and OVX+E_2_ mice (Suppl. Fig 2, H-J), further showing the complex interactions among inflammation and lipid metabolism in *Ifnγ*^−/−BM^ recipient mice. Together, these findings suggest that the ability to limit lipid peroxidation was preserved in young mice but impaired with aging, resulting in increased 4-HNE and its protein adducts when hepatic lipid metabolism was shifted toward fatty acid oxidation by menopausal E_2_ treatment.

### Menopausal E_2_ treatment induces genes associated with persistent inflammation in aging mice

To further investigate the underlying mechanism for the persistent inflammation associated with menopausal E_2_ treatment in aging mice, we performed bulk RNA-sequencing of liver tissues from sham and OVX+E_2_ treated aging mice. Consistent with increases in circulating inflammatory cytokines, hepatic transcriptomic analysis demonstrated coordinated induction of inflammatory and interferon-responsive genes following menopausal E_2_ treatment. 330 genes were differentially regulated between the two groups (Fig. 7A, Padj<0.05 and |log2 fold change| >log2(1.5)), indicating a distinct transcriptional profile in OVX+E_2_ mice compared with sham-operated mice. Among the upregulated genes in OVX+E_2_ mice, *Ltf*, *Ngp*, and *Elane* were associated with inflammation and myeloid activation. Hallmark gene-set enrichment analysis (GSEA) identified 12 significantly enriched gene sets (Fig. 7B). Inflammatory response, IFNγ response, IFNα response, and IL6-JAK-STAT3 showed positive enrichment in OVX+E_2_ mice. Complement activation and epithelial-mesenchymal transition pathways were also enriched, further supporting a pro-inflammatory and tissue-remodeling phenotype.

**Figure 7.**
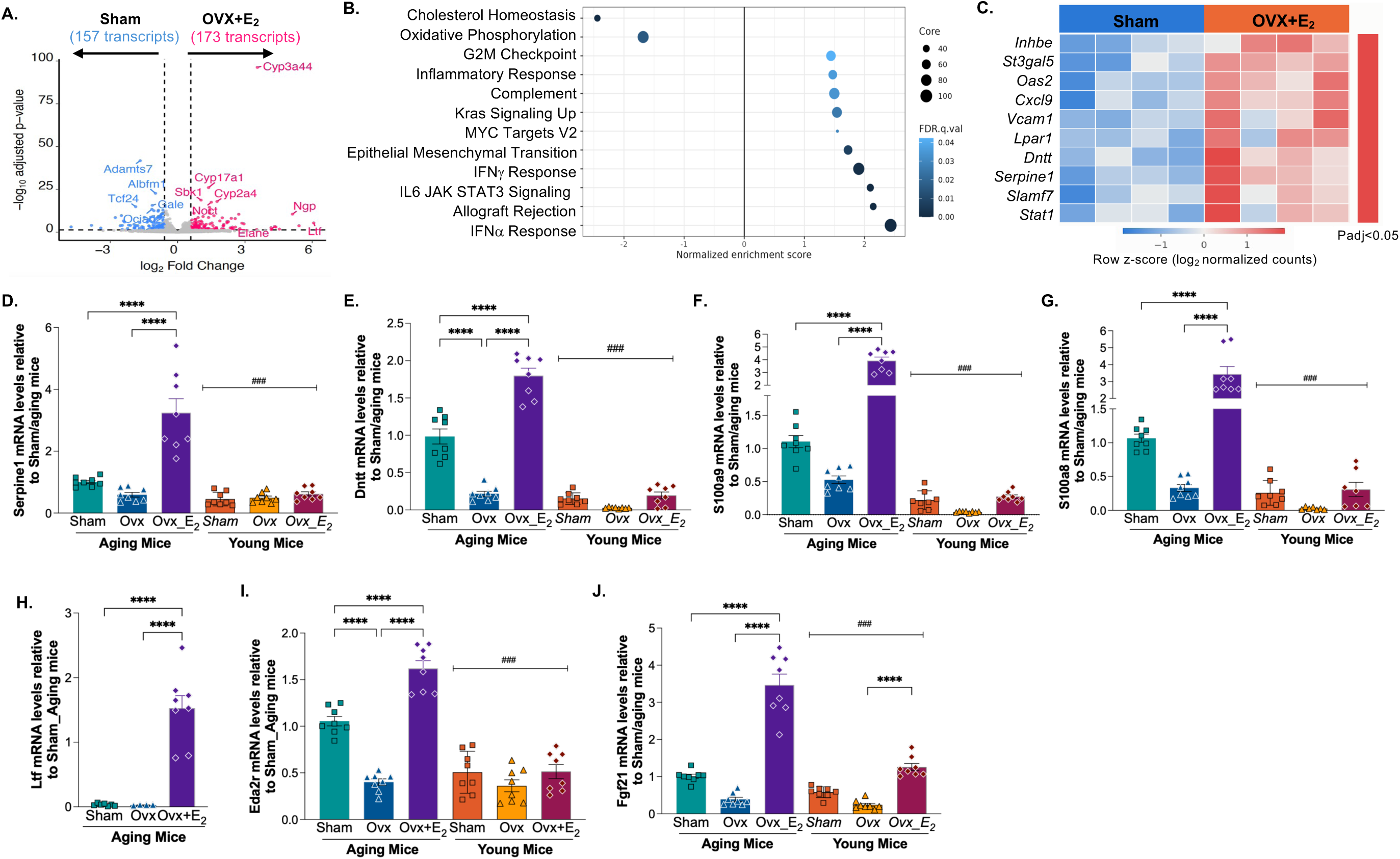
Liver transcriptomic profiling reveals enhanced inflammatory signaling in aging *Ldlr^−/−^* mice following OVX+E_2_ treatment. **A.** Volcano plot showing differential gene expression in aging OVX+E_2_ mice relative to aging sham controls. Positive log2 fold changes indicate higher expression in OVX+E_2_ mice. Highlighted genes meet adjusted *P*<0.05 and |log2 fold change| > log2(1.5). **B.** Hallmark gene sets meet Padj<0.05. Positive normalized enrichment scores indicate enrichment toward OVX+E_2_, whereas negative scores indicate enrichment toward sham. Point color indicates the reported FDR q value; point size indicates number of leading-edge genes. **C.** Expression of leading-edge genes contribute to inflammatory/interferon hallmark pathways shown in B. Colors show row z-scores of normalized counts. Leading-edge genes for selected pathways shown in Suppl. Fig. 4A. **D-J.** Comparison of hepatic expression of *Serpine1* (D), *Dntt* (E), *S100a9* (F), *S100a8* (G), *Ltf* (H*), Eda2r* (I), and *Fgf21* (J) between aging and young mice. Gene expression was normalized to 18s rRNA and is presented as fold change relative to the corresponding sham group in the aging cohort. Data are shown as mean ± SEM (n=7-8/group). Statistical significance was determined by two-way ANOVA with Sidak’s post hoc multiple-comparison tests. ^###^, *P*<0.001 between the aging and young cohort; ****, P<0.0001.

Leading-edge genes contributing to the enrichment of inflammatory and interferon response included *Cxcl9, Oas2, Stat1, Vcam1, Serpine1*, and *Slamf7* (Fig. 7C, Suppl. Fig. 4A). *Serpine1* encodes plasminogen activator inhibitor -1 (PAI-1), which is associated with inflammation, tissue remodeling, and fibrosis^40,41^. The effects of hormone therapy on circulating PAI-1 in humans has been inconsistent^42,43^. qPCR analyses supported increased hepatic *Serpine1* expression in aging OVX+E_2_ mice, whereas this response was attenuated in young OVX+E_2_ mice (Fig. 7D). Similar age-dependent expression patterns were observed for *Dntt, Cxcl9*, and *Stat1* (Fig. 7E, Suppl. Fig. 4, B-C). *Dntt* encodes terminal deoxynucleotidyl transferase (TdT), which is predominantly expressed in immature T- and B-cell precursors during V(D)J recombination^44^. Increased expression *of Dntt, Ltf, Ngp, and Elane,* suggests alterations in the hepatic immune-cell compartment accompanying the persistent inflammatory response to menopausal E_2_ treatment in aging mice.

Leading-edge analyses further demonstrated increased expression of myeloid-associated genes, including *S100a9* and *Ltf,* in response to E_2_ treatment in aging mice (Suppl. Fig. 4D). S100A9 along with S100A8, is almost exclusively expressed in neutrophils, monocytes, and myeloid cells^45^. Expression of these genes are upregulated upon inflammation and myeloid activation^45^. Extracellular S100A8/S100A9 can also function as a DAMPs (damage-associated molecular patterns) that activate Toll Like Receptor 4-dependent inflammatory signaling^45,46^. qPCR analyses further demonstrated that *S100a8* and *S100a9* expression was generally higher in the aging cohort than in the young cohort. In the aging cohort, their expression was higher in OVX+E_2_ mice than in sham and OVX mice, whereas differences between sham and OVX mice did not reach statistical significance (Fig. 7, F-G). *Ltf* encodes lactotransferrin (lactoferrin), an innate immune protein highly abundant in neutrophil granules and associated with innate immune responses^47,48^. *Ltf* was minimally expressed in the livers of aging sham and OVX mice but was markedly induced following E_2_ treatment (Fig. 7H). In contrast, *Ltf* transcripts were undetectable by qPCR in young mice, suggesting that *Ltf* induction reflected an age-dependent response rather than a general effect of E_2_ treatment. In aging *Ldlr^−/−^* mice reconstituted with *Ifnγ^− /-BM^*, induction of inflammatory transcripts contributing to interferon-responsive pathways was attenuated, whereas transcripts associated with myeloid-cell activation, such as *Ltf, S100a8,* and *S100s9*, remained elevated in OVX+E_2_ mice (Suppl. Fig. 5). Together, these findings suggest an enhanced pro-inflammatory signature characterized by myeloid cell activation in the aging liver following E_2_ treatment, whereas *Ifnγ^−/−BM^* transplantation suppressed downstream IFNγ-associated inflammatory signaling but could not fully prevent the aging- and E_2_-associated myeloid response.

In addition to inflammatory genes, gene ontology (GO) analysis identified lipid-related biological processes among differently expressed genes, including fatty acid metabolism, long-chain and unsaturated fatty acid metabolism, arachidonic acid and eicosanoid metabolism, and lipid hydroxylation (Suppl. Fig. 6, A-B). *Cyp3a44* encodes a P450 3A monooxygenase, annotated with lipid hydroxylation activity, along with additional cytochrome P450 genes, including *Cyp2c69* and *Cyp3A41a/16*, were upregulated in OVX+E_2_ mice (Fig. 7A and Suppl Fig. 6A). Among E_2_-responsive metabolic genes (Suppl. Fig. 6A), *Acot3* and *Acot4* encode acyl-CoA thioesterases that hydrolyze acyl-CoA to free fatty acid and CoA, providing substrates for fatty acid oxidation^49^. *Lipg* and *Pla2g4f* encode lipid-hydrolyzing enzymes that can release fatty acids from complex lipids and membrane phospholipids, thereby increasing substrate availability for downstream lipid metabolism^50,51^. Together with the increased pACC observed in aging OVX+E_2_ mice, these transcriptional changes indicate that menopausal E_2_ treatment promotes multiple pathways involved in fatty acid mobilization and oxidative metabolism.

Furthermore, transcriptomic analysis provided additional evidence of increased cellular and oxidative stress in aging OVX+E_2_ mice. *Cyp17a1* and *Sbk1* have been implicated in metabolic and cellular stress responses^52,53–55^. Both transcripts were upregulated by menopausal E_2_ treatment relative sham operation in the aging cohort (Fig. 7A). Ectodysplasin A2 receptor (*Eda2r)* is associated with cellular senescence and stress signaling^56^. *Eda2r* expression was increased in OVX+E_2_ mice compared with both sham and OVX mice in the aging cohort, whereas its expression remained low and largely unresponsive to hormone treatments in the young cohort (Fig. 7I). Among six differentially expressed genes annotated to Response to Oxidative Stress (GO: 0006979), five genes had higher expression in aging OVX+E_2_ mice, which was associated with enrichment in metabolic processes (Suppl. Fig. 6, B-C). Fibroblast growth factor 21 (FGF21) is a hepatokine that can be induced in response to metabolic and cellular stress^57^. qPCR analysis showed that *Fgf21* expression was increased in aging mice in response to menopausal E_2_ treatment, which response was attenuated in young mice (Fig. 7J). Collectively, these findings demonstrate that menopausal E_2_ treatment in aging mice induces a coordinated hepatic transcriptional program characterized by interferon and inflammatory signaling, an enhanced myeloid-associated signature, altered lipid metabolic pathways, and increased oxidative and cellular stress responses.

## Discussion

The present study investigated the mechanisms underlying the dissociation between the favorable metabolic effects of menopausal hormone therapy and its inconsistent cardiovascular benefits in mouse models of established atherosclerosis at young and relatively advanced ages. We demonstrate that, following lipid normalization, E_2_ treatment improved lipid metabolism in *Ldlr*^−/−^ mice at both ages and promoted hepatic pathways favoring fatty acid oxidation, as evidenced by increased phosphorylation of AMPKα and ACC and upregulation of PGC-1α. However, the ability to limit oxidative stress during this metabolic adaptation was age dependent. In young mice, E_2_ treatment reduced hepatic 4-HNE levels and 4-HNE protein adducts, suppressed systemic and plaque inflammation, preserved HDL antioxidant and cholesterol efflux functions, and reduced atherosclerotic burden to levels comparable to those in sham mice. In contrast, aging was associated with altered hepatic estrogen receptor expression, characterized by reduced *Esr1* and increased *Gper1* expression, alongside E_2_-induced *Nox1* expression and sustained elevations in hepatic 4-HNE and its protein adducts. Liver transcriptomic analyses further revealed that menopausal E_2_ treatment in aging mice enhanced pathways associated with fatty acid metabolism, lipid hydroxylation, and cellular stress responses, while concurrently activating interferon signaling, IL-6/JAK/STAT3 signaling, and innate immune pathways, together with an enhanced myeloid-associated inflammatory signature. Consistent with these findings, aging OVX+E_2_ mice exhibited persistently elevated circulating IFNγ, impaired HDL functions, and increased plaque inflammation and vulnerability despite improvements in hyperlipidemia. Suppression of hematopoietic IFNγ-mediated inflammation restored HDL function and the atheroprotective effects of E_2_ but did not fully attenuate the aging- and E_2_-associated myeloid inflammatory signature. Together, these findings support a model in which aging impairs the ability to constrain oxidative stress during E_2_-stimulated lipid metabolism, resulting in lipid peroxidation and persistent inflammatory signaling that compromise the cardiovascular benefits of E_2_ following OVX.

The observations reported in the present study parallel findings from major clinical trials of MHT, regarding the dissociation between its favorable metabolic effects and inconsistent cardiovascular benefits^5–7^. The “Timing Hypothesis” proposes that the cardiovascular benefits of MHT depend on when treatment is initiated relative to menopause, with limited or no benefit observed in women initiating MHT more than 10 years after menopause^10^. However, the independent contribution of chronological aging to cardiovascular responses to MHT remains unknown. The aging model in the present study consisted of 14-month-old *Ldlr*^−/−^ mice that had been exposed to a Western diet for three months, which may have accelerated metabolic and vascular aging. Although tissue responsiveness to sex hormones may decline with aging, it is not necessarily completely lost at this age^35,36^. Indeed, both young and aging mice exhibited improved lipid profile following E_2_ treatment, with ACC phosphorylation consistent with preserved hepatic pathways favoring fatty acid oxidation in OVX+E_2_ mice. However, only young mice demonstrated reduced plaque inflammation, improved HDL function, and regression of established atherosclerosis following E_2_ treatment. These findings suggest that the diminished cardiovascular efficacy of MHT with aging is not simply attributable to a generalized loss of estrogen responsiveness but may instead reflect age-dependent alterations in tissue-specific responses to E_2_ treatment.

Favorable hepatic metabolic responses to menopausal E_2_ treatment were attenuated with aging, potentially reflecting the marked reduction in hepatic ERα expression. Estrogens exert many of their metabolic benefits through ERα (*Esr1*), which regulates hepatic lipid metabolism, insulin sensitivity, and mitochondrial function. The aging-associated decline in hepatic ERα expression may contribute to the attenuated differences in hepatic TG accumulation between sham and OVX mice, as well as between OVX and OVX+E_2_, observed in the aging cohort compared with the young cohort. This is consistent with previous studies demonstrating that estrogens limit hepatic TG accumulation, in part by promoting fatty acid oxidation through hepatic ERα-dependent pathways^16,23^. Although estrogen retained metabolic activity in both age groups, the magnitude of these responses was greater in young mice. Specifically, compared with aging mice under the corresponding hormonal conditions, young mice exhibited greater reductions in circulating lipid levels, more favorable insulin homeostasis, and more pronounced decreases in hepatic TG accumulation in sham and OVX+E_2_ groups.

The marked reduction in hepatic *Esr1* expression may contribute to diminished metabolic responsiveness to estrogens with aging, whereas increased *Gper1* expression may alter estrogen signaling toward pathways associated with oxidative stress and inflammation. Hepatic *Gper1* expression was markedly elevated in the aging cohort relative to the young cohort, coinciding with enhanced hepatic oxidative stress signaling following E_2_ treatment. Previous studies have demonstrated that constitutive GPER activation can stimulate NOX1 activity and cause subsequent oxidative damage□ □. In this study, hepatic *Nox1* expression was selectively induced in the aging cohort, with higher expression in sham and OVX+E_2_ groups. This was associated with increased expression of genes involved in fatty acid metabolism and oxidation, including members of the *Cyp3a* family, *Acot3*, and *Acot4*, as well as enrichment of lipid hydroxylation and related GO terms. Transcriptomic analyses also identified enrichment of pathways associated with cellular responses to oxidative stress in aging E_2_-treated mice. For example, increased *Serpine*1 expression is not only associated with inflammation but also indicates increased cellular stress and impaired fibrinolytic activity^40,41^. These alterations accompanied by 4-HNE accumulation support oxidative injury in aging OVX+E_2_ mice. Hallmark gene set enrichment analysis identified negative enrichment of Oxidative Phosphorylation, driven by a limited subset of leading-edge genes. This finding may suggest altered mitochondrial energy metabolism, potentially associated with sustained oxidative stress during E_2_ treatment in aging mice. Notably, elevated liver 4-HNE levels in aging OVX mice may arise through lipid accumulation-associated lipotoxic stress, a mechanism distinct from that observed in aging OVX+E_2_ mice. The overall elevated liver 4-HNE levels in the aging cohort further reflex the association between hepatic lipid accumulation and age-dependent metabolic dysfunction. Thus, age-associated estrogen receptor remodeling and altered oxidative metabolic pathways compromises the ability to limit oxidative stress while E_2_ treatment improves liver accumulation.

Liver ERβ expression was increased with aging. ERβ exhibits a distinct cell type-specific distribution within the liver^58^. Whereas ERα is predominantly expressed in hepatocytes, ERβ is enriched in hepatic stellate cells, the principal fibrogenic cell population. Anti-fibrotic actions of estrogen are mediated primarily through ERβ rather than ERα or GPER^59,60^. Therefore, the increased ERβ expression observed in aging mice may represent an adaptive response to age-associated oxidative stress and tissue remodeling, potentially reflecting activation or expansion of hepatic stellate cells.

Beyond the potential pro-inflammatory effects of increased lipid peroxidation and oxidative stress, aging markedly influenced inflammatory responses to menopausal E_2_ treatment. Previous in vitro and in vivo studies have demonstrated that estrogens activate innate immune responses^61,62^. Recent evidence further demonstrates that estrogens promote NF-κB-mediated CXCL1 production through GPER1-dependent pathways, promoting neutrophil recruitment and extracellular trap formation^63^. However, how aging influences estrogen-mediated regulation of inflammation remains poorly understood. In the present study, the enrichment of inflammatory response, IFNα and IFNγ responses, IL-6/JAK/STAT3 signaling, and complement pathways in aging OVX+E_2_ mice compared with sham controls, were accompanied by increased expression of genes associated with inflammatory signaling, cellular stress responses, and immune cell activation. Notably, E_2_-induced upregulation of these genes including *Serpine1, Dntt, S100a9/8, Cxcl9, Stat1,* and *Ltf*, was more pronounced in aging mice than in young mice, suggesting that aging amplifies inflammatory responses to E_2_ treatment in the liver. Among these genes, *S100a8* and *S100a9* encode damage-associated molecular pattern proteins involved in myeloid cell activation and amplification of inflammatory signaling^45,46^, whereas *Cxcl9* encodes an IFNγ-responsive chemokine involved in immune cell recruitment. In contrast to aging mice, young mice exhibited coordinated improvements in both metabolic and cardiovascular outcomes following E_2_ treatment. Although several inflammatory genes that were markedly upregulated by E_2_ in aging mice also remained estrogen-responsive in young mice, their expression levels remained comparable to or below baseline levels (data not shown). Furthermore, OVX in young mice reduced circulating concentrations of several anti-inflammatory and proinflammatory cytokines, including IL-10, IL-4, IFNγ, IL-1β, and IL-6, following lipid normalization. However, OVX-induced circulating keratinocyte-derived chemokine (KC) and monocyte chemoattractant protein-1 (MCP-1) may enhance chemokine signaling associated with leukocyte recruitment. Notably, E_2_ treatment significantly reduced MCP-1 while partially restoring IL-10 concentrations in young mice. These findings raise the possibility that estrogen-induced inflammatory signaling represents a female-specific physiological response that is more effectively resolved in young animals but becomes exaggerated with aging. Whether these age-dependent differences reflect diminished immune resilience, impaired antioxidant defenses, mitochondrial dysfunction, or defective resolution of inflammation remains to be determined.

In summary, the present study demonstrates that aging fundamentally alters the biological response to menopausal estradiol treatment, in part through estrogen receptor remodeling characterized by reduced ERα dominance and increased GPER signaling, resulting in persistent inflammation. Persistent inflammation impairs HDL function and limits regression of established atherosclerosis despite preserved improvements in lipid metabolism and insulin homeostasis. Suppression of inflammation restores the cardiovascular benefits of E_2_, supporting inflammation-driven, non-lipid mechanisms as major contributors to residual ASCVD risk during hormone therapy. These findings provide a mechanistic insight into the age-dependent loss of cardiovascular benefits of hormone therapy and identify hepatic estrogen receptor remodeling, oxidative stress, and inflammatory signaling as potential therapeutic targets for improving cardiovascular outcomes in postmenopausal women receiving hormone therapy.

## AUTHORSHIP CONTRIBUTION STATEMENT

CDL and CG performed the experiments, obtained data, and prepared the manuscript. BL, MZ, JR, JA, RB, AV, and SC performed experiments and obtained data. YW performed RNAseq analyses and edited the manuscript. SC, YX, MFL, and JMS analyzed the data and edited the manuscript. LZ developed the overall concept and study design, prepared the figures, wrote and edited the manuscript. All authors approved the final version.

## ACKNOWLEDGMENTS

The authors acknowledge the helpful assistance of the Vanderbilt Hormone Assay Core (supported by NIH grant DK020593 to the Vanderbilt Diabetes Research Center). We also acknowledge excellent support of the Vanderbilt Mouse Metabolic Phenotyping Core (supported by NIH grant DK59637). We additionally recognize the support of the Vanderbilt Diabetes Research Center (P30DK020593) and the Vanderbilt Digestive Disease Research Center (P30DK058404).

---

**Supplemental Table 1:** TaqMan Assays used for qPCR.

| **Protein** | **Gene Symbol** | **Assay ID** |
| --- | --- | --- |
| RNA, 18S Ribosomal 1 | *18S* | Hs99999901_s1 |
| Estrogen Receptor 1 ($\alpha$) | *Esr1* | Mm00433149_m1 |
| Estrogen Receptor 2 ($\beta$) | *Esr2* | Mm00599821_m1 |
| G Protein-Coupled Estrogen Receptor 1 | *Gper1* | Mm02620446_s1 |
| NADPH Oxidase 1 | *Nox1* | Mm00627696_m1 |
| S100 Calcium Binding Protein A8 | *S100a8* | Mm00496696_g1 |
| S100 Calcium Binding Protein A9 | *S100a9* | Mm00656925_m1 |
| Plasminogen activator inhibitor-1 | *Serpine1* | Mm00435858_m1 |
| C-X-C Motif Chemokine Ligand 9 | *Cxcl9* | Mm00434946_m1 |
| Signal Transducer and Activator of Transcription 1 | *Stat1* | Mm01257286_m1 |
| Fibroblast Growth Factor 21 | *Fgf21* | Mm00840165_g1 |
| Ectodysplasin A2 Receptor | *Eda2r* | Mm00723601_m1 |
| Terminal deoxynucleotidyl transferase (TdT) | *Dntt* | Mm00493500_m1 |
| Lactotransferrin (lactoferrin) | *Ltf* | Mm00434787_m1 |

