## Supplemental figures for "The Cardiovascular Benefit of Menopausal Hormone Treatment is Age-Dependent in *Ldlr^−/−^* Mice"

### Supplemental Figure 1

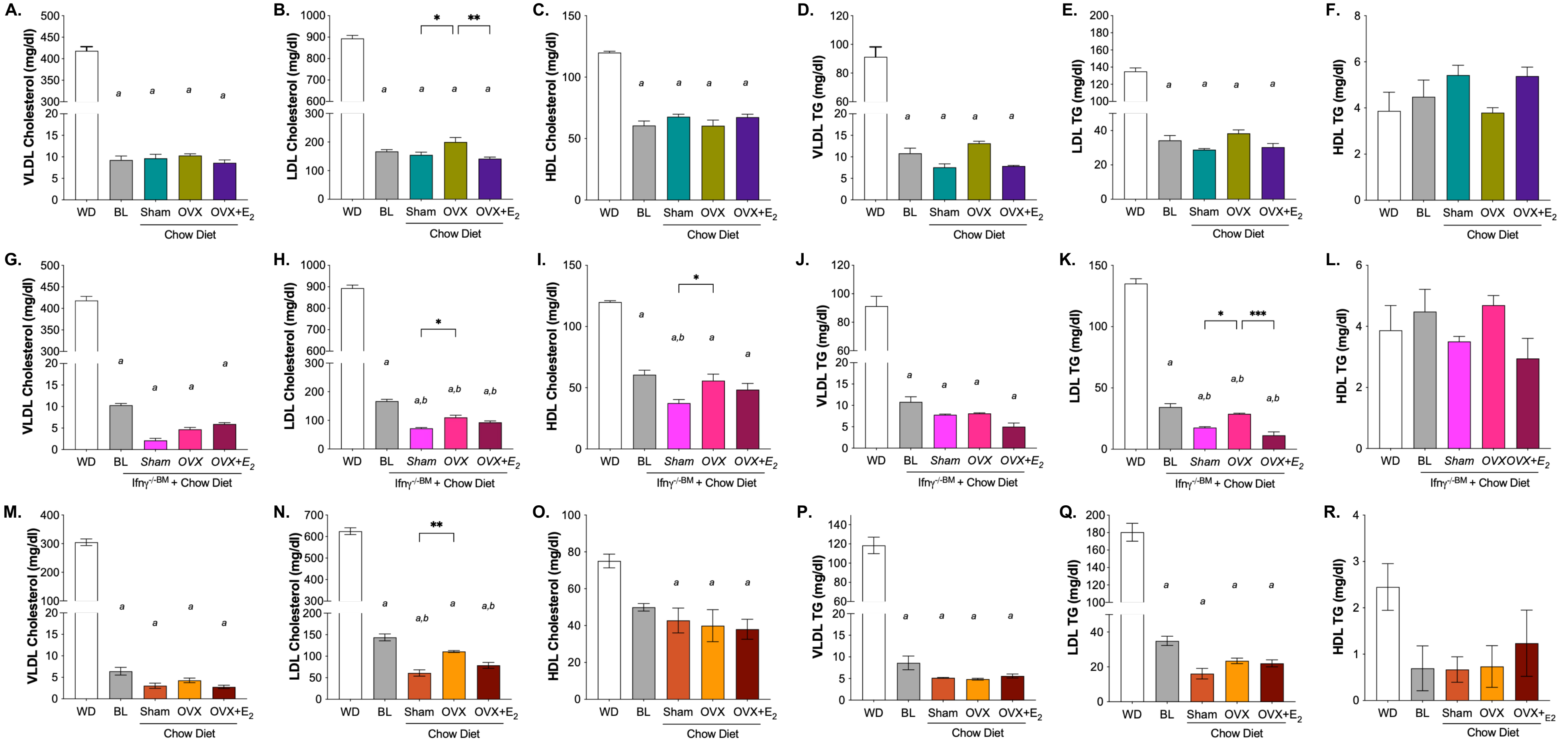


**Suppl. Fig 1. Serum cholesterol and triglycerides (TG) distribution across lipoproteins fractions.** Serum cholesterol and TG distribution across lipoprotein fractions in aging mice (A-F), *Ifnγ-/-BM* recipient mice (G-L), and young mice (M-R) across hormone treatment groups. FPLC analysis was performed using serum samples pooled from 2-3 mice. Data are presented as mean ± SEM (n>=7/group). Statistical analyses were performed using one-way ANOVA. *a*, *P*<0.05 compared with Western diet conditions (WD); *b*, P<0.05 compared with baseline (BL) *, P<0.05; **, P<0.01, ***, P<0.001.

#### Supplemental Figure 2

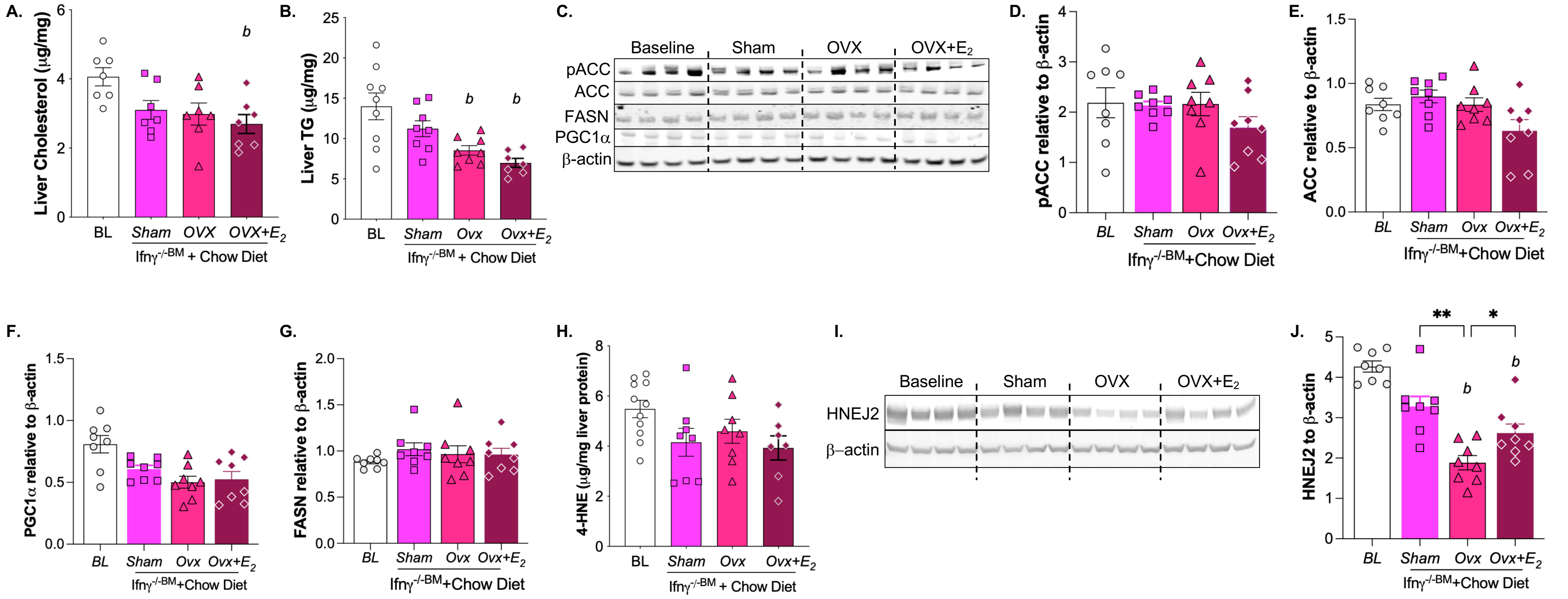


**Suppl. Fig 2. Hepatic lipid content and key proteins involved lipid metabolism in *Ifnγ$^{-/-BM}$* recipient mice. A-B.** Hepatic cholesterol and triglyceride (TG) contents. **C-G**. Representative immunoblots and quantification of pACC, ACC, FASN, and PGC1α. **H.** Hepatic 4-HNE content. **I-J.** Representative immunoblots and quantification of 4-HNE protein adducts detected using HNEJ2 antibody. Data are presented as mean ± SEM (n>=7/group). Statistical analyses were performed using one-way ANOVA with Tukey's multiple comparisons test. *b*, $P$<0.05 compared with baseline (BL) conditions. *, $P$<0.05; **, $P$<0.01.

#### Supplemental Figure 3

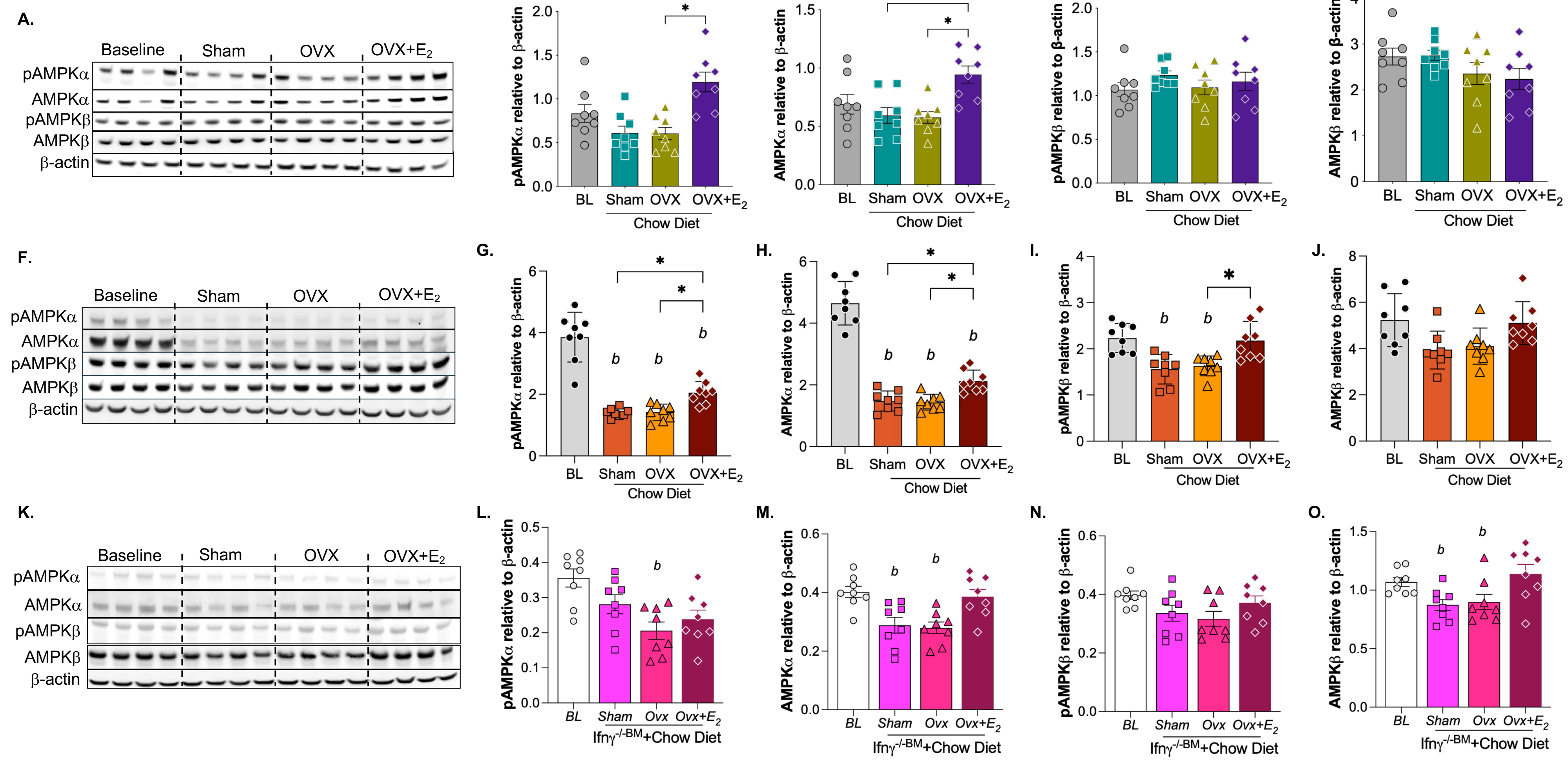


**Suppl. Fig 3. Hepatic AMPKα/β expression and phosphorylation across aging, young, and the *Ifnγ*$^{-/-BM}$ recipient mice.** Representative immunoblots quantifications of phosphorylated AMPKα/β (pAMPKα/β) and total AMPKα/β across aging **(A-E),** young **(F-J),** and the *Ifnγ*$^{-/-BM}$ recipient mice **(K-O).** Data are presented as mean ± SEM (n>=7/group). Statistical analyses were performed using one-way ANOVA with Tukey's multiple comparisons test. *b*, *P*<0.05 compared with baseline (BL) conditions. *, *P*<0.05.

#### Supplemental Figure 4

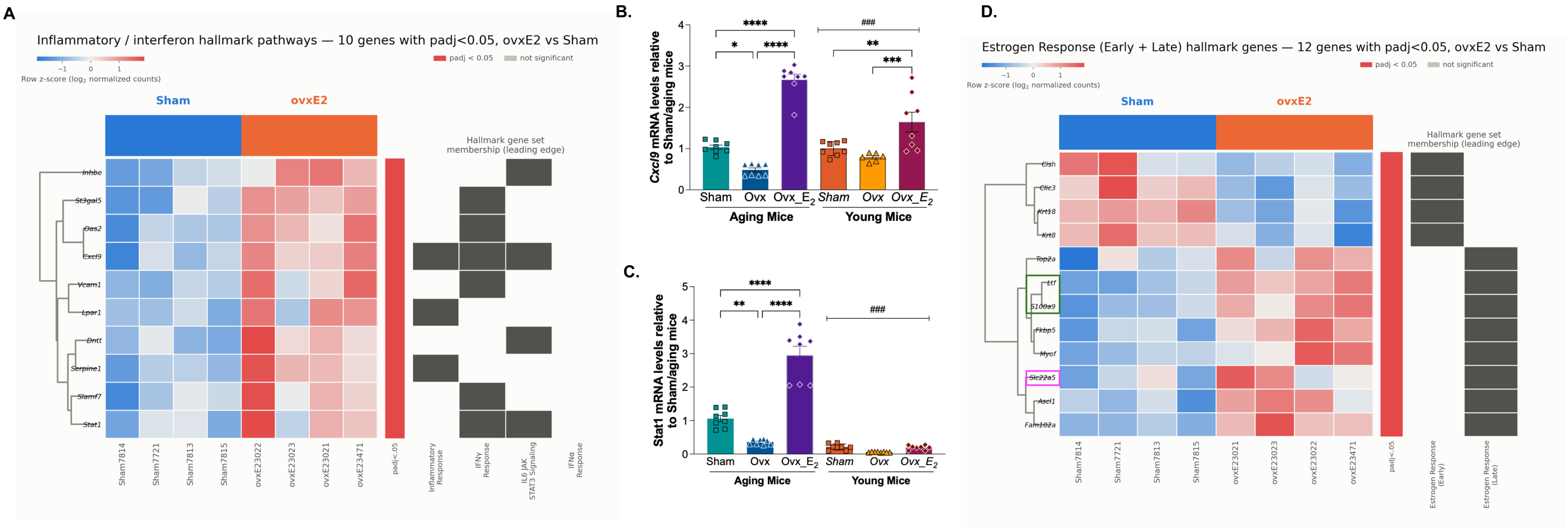


**Suppl. Fig. 4. Hepatic expression of inflammatory and estrogen-response gene sets. A.** Inflammatory and interferon Hallmark pathways. Heatmap showing 10 genes annotated to one or more of four Hallmark gene sets shown in Fig. 7B. Heatmaps display row-wise z-scores of normalized RNA-seq counts for significantly differentially expressed genes in liver from sham and OVX+$E_2$ mice. Each column represents an individual mouse. Differential expression was determined using DESeq2 with Wald test and Benjamini-Hochberg false discovery rate (FDR) correction; genes shown in each panel passed *padj* < 0.05. Red and blue indicate relatively higher and lower expression, respectively, based on row z-scores; gray boxes in the annotation matrices indicate gene-set membership. Gene set definitions were drawn from Hallmark leading-edge gene lists. **B-C.** qPCR validation of *Cxcl9* and *Stat1* expression in both aging and young cohorts. Gene expression was normalized to 18s rRNA. **D.** Differentially expressed genes belonging to the Hallmark Estrogen Response Early and Estrogen Response Late gene sets. *Ltf* and *S100a9*, outlined in green, are associated with myeloid/innate immune responses. *Slc22a5,* outlined in purple, encodes a plasma membrane carnitine transporter involved in cellular carnitine uptake and mitochondria long-chain fatty acid oxidation. Data are presented as fold change relative to the sham condition in the aging cohort. qPCR data are shown as mean ± SEM (n>=7/group). Statistical significance was determined using two-way ANOVA with Sidak's post hoc multiple-comparison tests. $^{###}$, *P*<0.001 between aging and young cohorts; *, *P*<0.05, **, P<0.01, ***, *P*<0.001, ****, P<0.0001.

#### Supplemental Figure 5

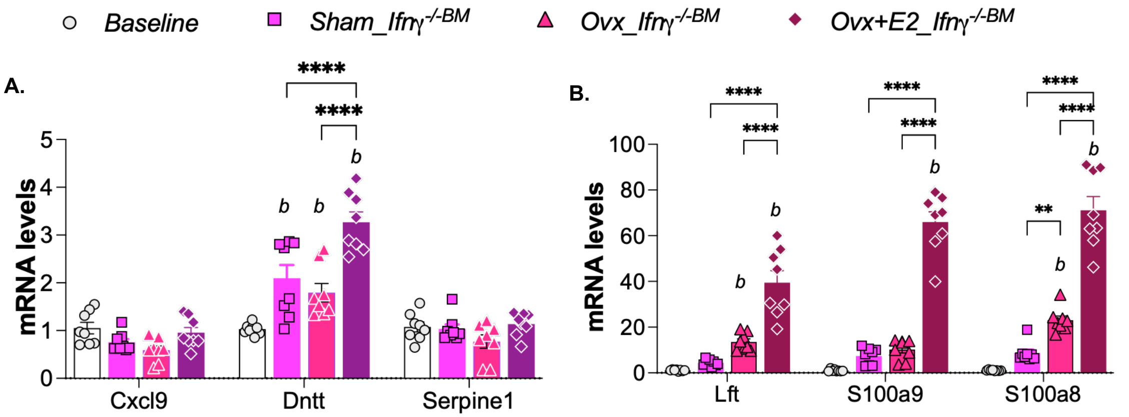


**Suppl. Fig 5. qPCR analyses of inflammatory and myeloid-associated gene expression in *Ifnγ*$^{/-BM}$ recipient mice. A.** $E_2$–induced upregulation of inflammatory gene, including Cxcl9 and Serpine1, observed in the aging cohort was attenuated in *Ifnγ*$^{/-BM}$ recipient mice. B. Expression of myeloid-associated genes remained elevated in OVX+$E_2$ mice despite of the *Ifnγ* $^{/-BM}$ transplantation. Data are presented as mean ± SEM (n>=7/group). Statistical significance was determined using two-way ANOVA with Sidak's post hoc multiple-comparison tests. *b*, *P*<0.05 compared with baseline (BL) conditions. ****, *P*<0.0001.

### Supplemental Figure 6

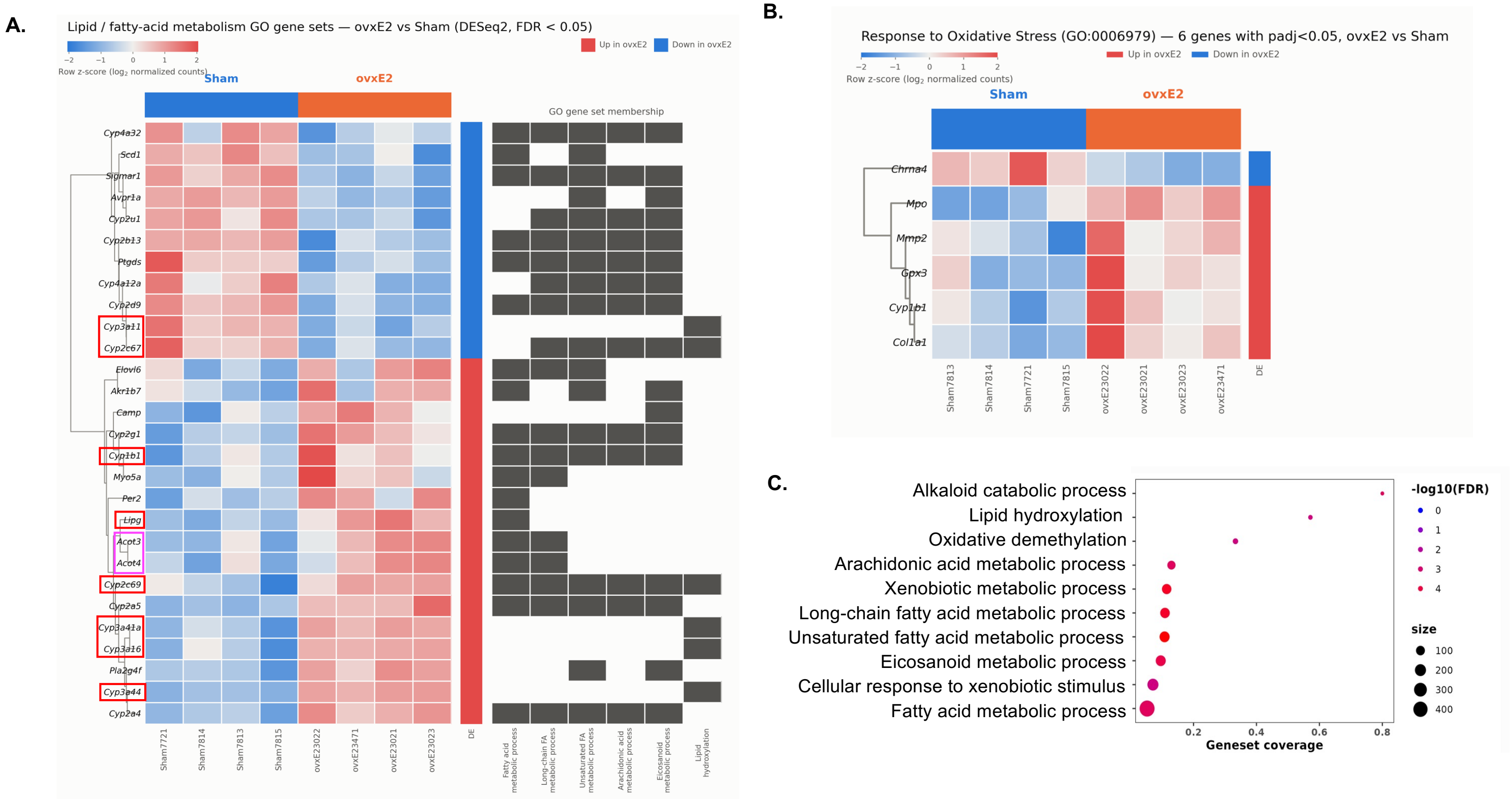


**Suppl. Fig. 6. Hepatic transcriptomic analysis identifies differentially expressed genes associated with lipid metabolism and oxidative stress.** Heatmaps show row-wise z-scores of normalized RNA-seq counts for significantly differentially expressed genes in livers from sham and Ovx+$E_2$ mice. Each column represents an individual mouse. **A.** Differentially expressed genes associated with lipid and fatty-acid metabolism Gene Ontology (GO) gene sets**.** The annotation matrix on the right indicates membership of each gene in the indicated GO pathways shown in panel B. The vertical annotation bar indicates genes upregulated (red) or downregulated (blue) in Ovx+$E_2$ relative to sham mice. Gene labels outlined in red mark genes in Lipid Hydroxylation or Response to Oxidative Stress gene sets, whereas labels outlined in purple indicate genes encoding acyl-CoA thioesterases, proteins with lipase activity that may increase fatty acid availability for downstream metabolism. **B.** Differentially expressed genes belonging to Response to Oxidative Stress (GO:0006979). Differential expression was determined by DESeq2 with Wald test, Benjamini-Hochberg FDR correction; genes shown in each panel passed *padj* < 0.05. Red and blue in the heatmaps indicate relatively higher and lower expression, respectively, based on row z-scores. Gene set definitions were obtained from MSigDB Gene Ontology Biologic Process terms. **C.** Six out of top 10 significantly enriched biological processes (GO) are associated with lipid metabolism.
